# The stem cell-specific protein TRIM71 inhibits maturation and activity of the pro-differentiation miRNA let-7 via two independent molecular mechanisms

**DOI:** 10.1101/2021.01.29.428807

**Authors:** Lucia A. Torres-Fernández, Sibylle Mitschka, Thomas Ulas, Kilian Dahm, Matthias Becker, Kristian Händler, Julia Windhausen, Joachim L. Schultze, Waldemar Kolanus

**Affiliations:** Life and Medical Sciences Institute (LIMES), Molecular Immunology and Cell Biology, University of Bonn, 53115 Bonn, Germany; Systems Medicine, German Center for Neurodegenerative Diseases (DZNE), 53175 Bonn, Germany; Platform for Single Cell Genomics and Epigenomics (PRECISE), German Center for Neurodegenerative Diseases (DZNE), 53175 Bonn, Germany; Life and Medical Sciences Institute (LIMES), Genomics and Immunoregulation, University of Bonn, 53115 Bonn, Germany

**Author notes:** Equal contribution (Lucia A. Torres-Fernández, Sibylle Mitschka, Thomas Ulas).

**Keywords:** TRIM71/LIN-41, RNA-binding protein (RBP), E3 ubiquitin ligase, miRNAs, let-7 family, AGO2, LIN28, TUT4, embryonic development, hepatocellular carcinoma, stem cells and cancer biology

## Abstract

The stem cell-specific RNA-binding protein TRIM71/LIN-41 was the first identified target of the pro-differentiation and tumor suppressor miRNA let-7. TRIM71 has essential functions in embryonic development and a proposed oncogenic role in several cancer types, such as hepatocellular carcinoma. Here, we show that TRIM71 regulates let-7 expression and activity via two independent mechanisms. On the one hand, TRIM71 enhances pre-let-7 degradation through its direct interaction with LIN28 and TUT4, thereby inhibiting let-7 maturation and indirectly promoting the stabilization of let-7 targets. On the other hand, TRIM71 represses the activity of mature let-7 via its RNA-dependent interaction with the RNA-Induced Silencing Complex (RISC) effector protein AGO2. We found that TRIM71 directly binds and stabilizes let-7 targets, suggesting that let-7 activity inhibition occurs on active RISCs. MiRNA enrichment analysis of several transcriptomic datasets from mouse embryonic stem cells and human hepatocellular carcinoma cells suggests that these let-7 regulatory mechanisms shape transcriptomic changes during developmental and oncogenic processes. Altogether, our work reveals a novel role for TRIM71 as a miRNA repressor and sheds light on the precise mechanistic dual regulation of let-7.

## Introduction

TRIM71/LIN-41 was first discovered as a heterochronic gene in the nematode *C. elegans*, where it was identified as a major target of the pro-differentiation lethal 7 (let-7) miRNA family^1,2^. Later studies confirmed the conservation of TRIM71/LIN-41 and its repression by let-7 miRNAs in other species^2–7^, highlighting the importance of this TRIM-NHL protein in the control of developmental processes. In mice, TRIM71 is highly expressed during early embryonic development and its expression starts to decrease at around E10.5 due to a progressive increase of let-7 and miR-125 miRNAs in the course of differentiation^5^. Despite its short expression window, TRIM71 is essential for embryonic development in several vertebrate and invertebrate species^8^. Embryonic lethality in mice occurs at E12.5 and is accompanied by neural tube closure defects^9–11^, underscoring a fundamental role of TRIM71 in the development of the nervous system. Furthermore, recurrent TRIM71 mutations have been identified in congenital hydrocephalus (CH) patients^12^, a brain developmental disease which is associated with neural tube closure defects^13^, and is characterized by an abnormal accumulation of cerebrospinal fluid (CSF) within the brain ventricles. Interestingly, TRIM71 expression has been observed in CSF-producing ependymal cells in the adult brain^10^, revealing yet-unknown post-natal TRIM71 functions in the nervous system. Last, TRIM71 is known to be expressed in adult mouse testes^14^, and has been recently found to play an essential role in the embryonic development of the germline as well as in adult spermatogenesis^15^.

On the molecular level, TRIM71 has a set of specialized domains enabling its function as an E3 ubiquitin ligase^16–19^ and an mRNA-binding and repressor protein^11,20–24^. A number of protein and mRNA targets regulated by TRIM71 have been linked to the developmental phenotypes observed in TRIM71-deficient mice^11,16,18,20–22,24,25^. TRIM71 has also been described as a miRNA-binding protein^26^. Furthermore, several proteins involved in the miRNA pathway, namely AGO1, AGO2, AGO4, DICER and LIN28B, are confirmed protein interactors of TRIM71^14,20,21,24,27^. However, the role of TRIM71 in the regulation of miRNAs remains poorly understood. Previous studies in our lab showed that TRIM71-deficient murine embryonic stem cells (ESCs) had an altered miRNA expression landscape as compared to wild type ESCs^11^, but it is yet unclear how TRIM71 regulates miRNA expression. Furthermore, TRIM71 was reported to decrease global miRNA activity via ubiquitylation and proteasomal degradation of AGO2^14^. However, TRIM71-mediated changes in AGO2 stability were not observed in several other studies^11,16,20,21,24^. Thus, the functional significance of the interaction between TRIM71 and AGO2 remains elusive.

We therefore aimed to investigate the role of TRIM71 in the regulation of miRNA expression and activity. Our results show that TRIM71 is not only a let-7 target, but also a let-7 repressor which employs two independent mechanisms to regulate let-7 expression and activity. For the regulation of let-7 expression, TRIM71 interacts with the let-7 repressor complex formed by LIN28 and TUT4, which intercepts let-7 miRNAs at their precursor stage and labels them for degradation. Furthermore, the interaction between TRIM71 and AGO2 results in a specific repression of let-7 activity, without either affecting AGO2 stability or globally inhibiting miRNA activity. Together, these TRIM71-mediated miRNA repression mechanisms contribute to regulate the transcriptomic landscape of ESCs and hepatocellular carcinoma cells, leading to the stabilization of multiple let-7 targets during developmental and oncogenic processes.

## Results

### TRIM71 interferes with the last processing step of let-7 biogenesis in ESCs

TRIM-NHL proteins have been connected to the miRNA pathway in several species^28^. In a previous study, we analyzed the impact of TRIM71 deficiency on the transcriptome of murine ESCs by performing RNA sequencing in undifferentiated wild type (WT, *Trim71^fl/fl^*) and *Trim71* knockout (KO, *Trim71^-/-^*) ESCs^11^. Reanalysis of these data focusing on miRNA expression shows that the miRNome of TRIM71-deficient ESCs is substantially altered, with increased expression of differentiation-promoting brain-specific and gonad-specific miRNAs and reduced expression of several ESC-specific miRNAs (Fig. 1A and ^11^). Interestingly, all members of the Let-7 miRNA family were found upregulated in TRIM71-deficient ESCs (Fig. 1B). qRT-PCR analysis confirmed a significant two-fold upregulation of let-7a and let-7g species and the downregulation of the stem cell-specific miR-302 in TRIM71-deficient ESCs (Fig. 1C and Suppl. Fig. 1A-D). Other miRNAs such as miR-125, which targets *Trim71* mRNA for degradation^5,9^, or the stem cell-specific miR-294, which functionally counteracts let-7 miRNAs^29^, remained unaltered upon TRIM71 depletion (Fig. 1C and Suppl. Fig. 1E-H).

**Figure 1.**
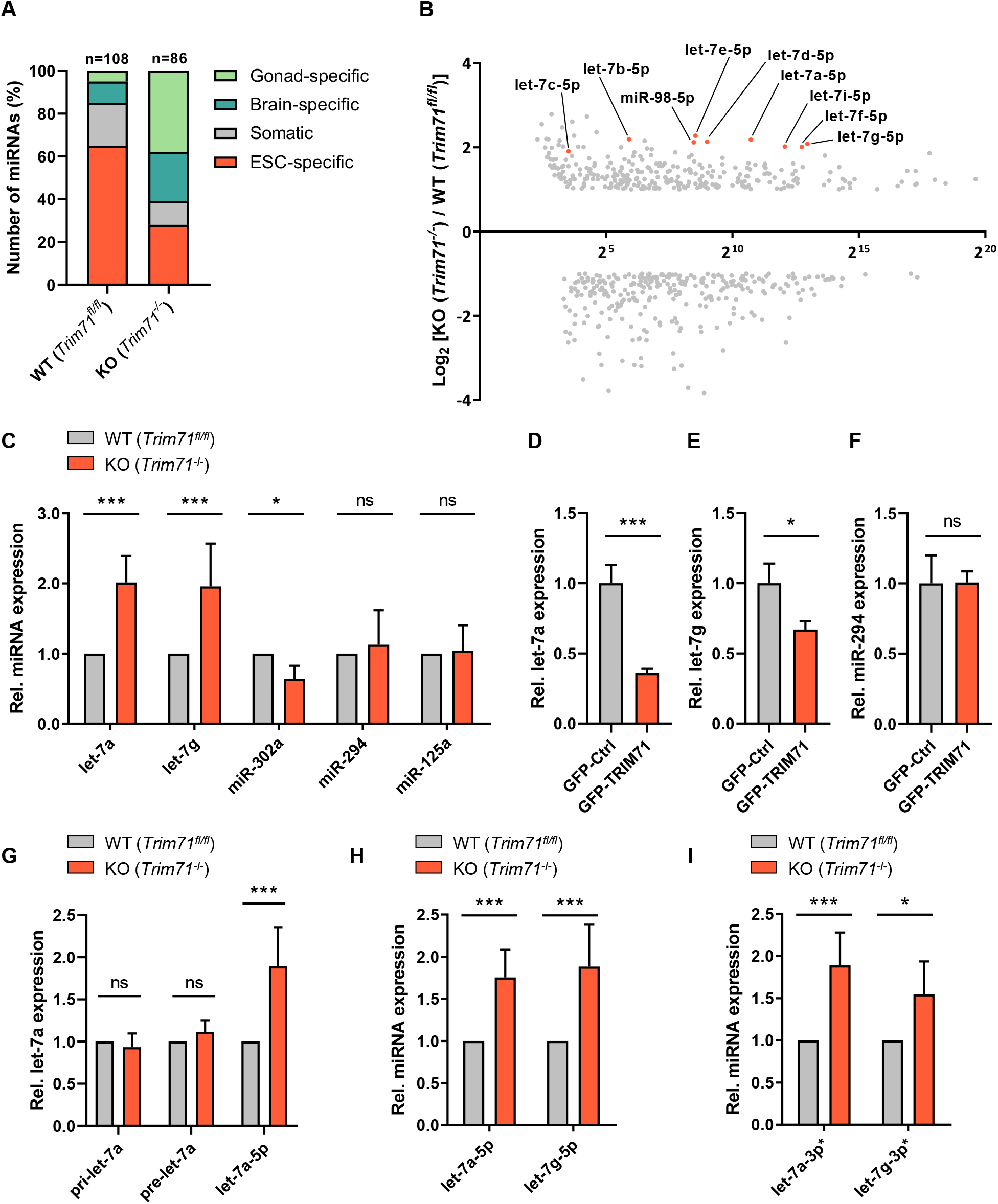
TRIM71 interferes with the last processing step of let-7 biogenesis in ESCs. **A)** Representation of miRNome differences observed in miRNASeq data obtained from wild type (WT, *Trim71^fl/fl^*) and *Trim71* knockout (KO, *Trim71^-/-^*) ESCs (GSE62509)^11^. MiRNAs were classified as previously described^65^ into four categories (gonad-specific, brain-specific, ESC-specific or somatic miRNAs) and displayed as percentages (n=590). **B)** Scatterplot showing each miRNA fold change (FC) as Log_2_(KO/WT), versus each miRNA abundance in WT ESCs (GSE62509)^11^. Guide strands (5p) of all let-7 family members are marked. **C)** RT-qPCR showing relative expression of the indicated miRNAs in WT and *Trim71* KO ESCs (n=3-10). **D)** RT-qPCR showing relative expression of let-7a, **E)** let-7g and **F)** miR-294 miRNAs in WT ESCs transiently overexpressing GFP-Ctrl or GFP-TRIM71 48 hpt (n=3). **G)** RT-qPCR showing relative pri-, pre- and mature let-7a expression in WT and *Trim71* KO ESCs (n=3-5). **H)** RT-qPCR showing relative guide (5p) and **I)** passenger (3p*) strand levels of mature let-7a and let-7g miRNAs in in WT and *Trim71* KO ESCs (n=3-5). RT-qPCR quantification of all miRNAs was normalized to the levels of the housekeeping U6 snRNA. Error bars represent SD. ***P-value < 0.005, **P-value < 0.01, *P-value <0.05, ns = non-significant (unpaired student’s t-test). See also Suppl. Fig. 1-2.

*Trim71* mRNA is a well-known let-7 target^5,9,1,2,4,7,30,31^, but the fact that let-7 members are upregulated upon TRIM71 depletion suggests a negative feedback loop exerted by TRIM71 on let-7 miRNAs. To confirm that let-7 upregulation was a direct consequence of TRIM71 depletion, and not an artefact of clonal selection in *Trim71* KO ESCs, we induced the deletion of the floxed *Trim71* alleles in WT (Trim71^fl/fl^/Rosa26-CerER^T2^) ESCs via 4-hydroxytamoxifen (4-OHT) treatment and measured let-7a levels over time. We detected a progressive increase of let-7a miRNA expression correlated with the loss of *Trim71* mRNA and protein upon 4-OHT treatment (Suppl. Fig. 2A-B). Conversely, TRIM71 overexpression in WT ESCs resulted in a significant downregulation of let-7a and let7g, but not miR-294 (Fig. 1D-F). These results revealed a specific and direct involvement of TRIM71 in the regulation of let-7 expression in ESCs.

To determine at which stage of let-7 biogenesis this regulation occurs, we measured primary (pri-let-7a), precursor (pre-let-7a), and mature let-7a levels in WT and *Trim71* KO ESCs. We found that mature let-7a levels were significantly upregulated in *Trim71* KO ESCs, whereas pri- and pre-miRNAs remained unaltered (Fig. 1G). This indicated that TRIM71 affects either the last step of let-7 maturation (i.e. pre-let-7 processing to mature let-7 miRNA duplex), or the stability of the mature miRNA (i. e. half-life of the let-7 guide strand). In order to distinguish between these two possibilities, we analyzed the relative abundance of guide (5p) and passenger (3p*) strands of let-7a and let-7g miRNAs in WT and *Trim71* KO ESCs and found both strands equally increased in *Trim71* KO ESCs (Fig. 1H-I and Suppl. Fig. 2C-D). This suggested that TRIM71 interferes with the last processing step of let-7 biogenesis, resulting in the downregulation of the mature miRNA duplex.

### TRIM71 depends on LIN28A for the downregulation of let-7 expression in ESCs

LIN28 proteins have an established role as inhibitors of let-7 biogenesis in undifferentiated stem cells^32,33^. Two LIN28 protein isoforms exist – LIN28A and LIN28B – and both isoforms are able to directly interact with pre-let-7 to promote its degradation^34,35^. Additionally, LIN28B sequesters pri-let-7 and pre-let-7 inside the nucleus, preventing its processing and nuclear export^36^. Notably, TRIM71 and LIN28 proteins have highly similar expression patterns during development in *C. elegans*, zebrafish and mouse^37^. Furthermore, their expression is highly correlated in healthy adult human tissues (Suppl. Fig. 3A-B), as well as in tumor samples of different cancer types (Suppl. Fig. 3B-C) according to the data available at the Gene Expression Profiling Interactive Analysis (GEPIA) server^38^.

In order to investigate whether TRIM71-mediated let-7 regulation is linked to LIN28 function, we generated *Lin28a* KO ESCs (*Trim71^fl/fl^; Lin28^−/−^*) as well as double *Trim71/Lin28a* KO ESCs (*Trim71^−/−^; Lin28^−/−^*) (Fig. 2A-C). Similar to what we previously reported for *Trim71* KO ESCs^11^, and agreeing with previous studies in *Lin28a* KO ESCs^39^, we found that stemness was unimpaired in all KO cell lines, as assessed by an unaltered ESC-characteristic morphology (Suppl. Fig. 4A-B) and a robust expression of stemness markers such as *Nanog, Myc* and *Pou5f1/Oct4* in all KO ESC lines (Suppl. Fig. 4C-E). We then measured the relative levels of pri-let-7a, pre-let-7a and mature let-7a-5p and let-7g-5p miRNAs in WT, *Trim71* KO, *Lin28a* KO and double KO ESCs. Again, we found no changes in either primary or precursor miRNA expression (Fig. 2D-E), while the levels of mature let-7a-5p and let-7g-5p – but not miR-294 (Suppl. Fig. 4F) – were significantly increased in all KO cell lines (Fig. 2F-G). Of note, the upregulation of let-7 was not indirectly caused by changes in the expression of *Lin28b* or *Zchcc11/tut4* (Suppl. Fig. 4G-H), which are also known to participate in the repression of let-7 expression^33,35,40^.

**Figure 2.**
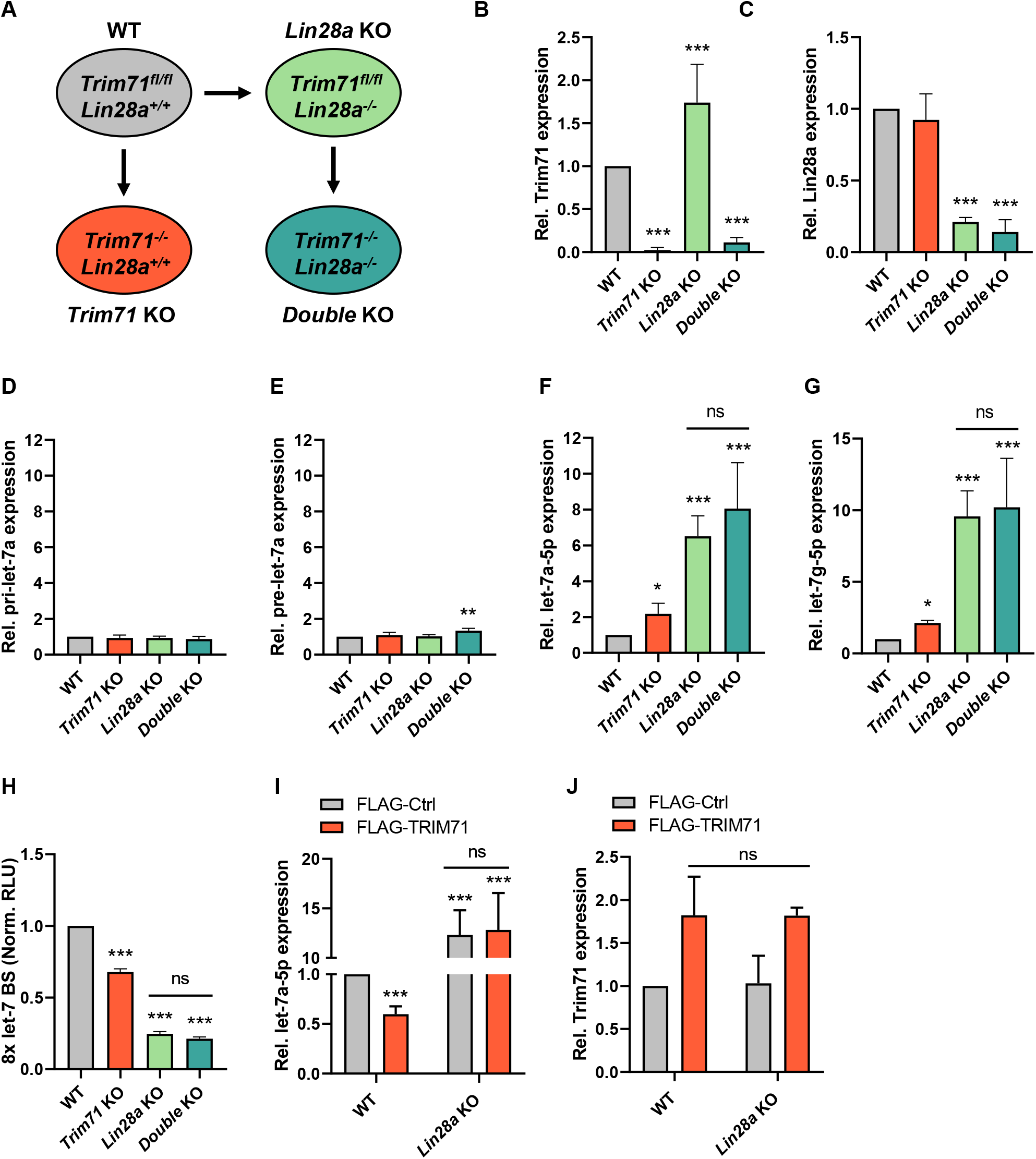
TRIM71 depends on LIN28A for the downregulation of let-7 expression in ESCs. **A)** Schematic representation for the generation of the indicated ESC lines. *Lin28a* KO ESCs were generated from WT (*Trim71^fl/fl^*) ESCs via TALENs. *Trim71* KO and *Trim71-Lin28a* double KO were generated by addition of 4-OHT to WT and *Lin28a* KO ESCs, respectively (see Methods for details). **B)** RT-qPCR showing relative mRNA expression of *Trim71* and **C)** *Lin28a* in the generated ESC lines (n=6-8). **D)** RT-qPCR showing relative expression of pri-let-7a, **E)** pre-let-7a, **F)** mature let-7a-5p and **G)** mature let-7g-5p miRNAs in the different ESC lines (n=4-8). **H)** Let-7 reporter assay upon transient transfection of a Renilla Luciferase reporter under the control of a 3’UTR containing 8x Let-7 binding sites (BS) in the different ESC lines (n=3). Norm. RLU = Normalized Relative Light Units. **I)** RT-qPCR showing relative expression of mature let-7a-5p and **J)** *Trim71* mRNA in WT and *Lin28a* KO ESCs transiently overexpressing FLAG-Ctrl or FLAG-TRIM71 48 hpt (n=3-6). RT-qPCR quantification of miRNAs and mRNAs was normalized to the levels of the housekeeping U6 snRNA and *Hprt* mRNA, respectively. Error bars represent SD. ***P-value < 0.005, **P-value < 0.01, *P-value <0.05, ns = non-significant (unpaired student’s t-test between WT and each KO condition, unless indicated by a line joining the two compared conditions). See also Suppl. Fig. 4.

Importantly, although the upregulation of let-7 was stronger in *Lin28a* KO ESCs than in *Trim71* KO ESCs, no additional increase of mature let-7a/g was observed for the double KO compared to the single *Lin28a* KO (Fig. 2F-G). These results were confirmed by let-7 reporter assays in ESCs, in which a luciferase reporter containing 8x let-7 binding sites (BS)^41^ was significantly repressed in all KO lines, but no additional repression was observed in the double KO as compared to *Lin28a* KO ESCs (Fig. 2H). This suggested that the suppressive effect of TRIM71 on let-7 expression in ESCs is dependent on LIN28A. Indeed, the overexpression of TRIM71 in *Lin28a* KO ESCs had no effect on let-7 expression, while let-7a levels were reduced by about 40% upon TRIM71 overexpression in WT ESCs (Fig. 2I) despite TRIM71 overexpression levels being comparable in both cell lines (Fig. 2J). These results collectively showed that TRIM71-mediated let-7 downregulation in ESCs depends on LIN28A.

Our let-7 reporter assay (Fig. 2H) suggested that the changes in let-7 expression observed in *Trim71* KO, *Lin28a* KO and double KO ESCs may have an impact on let-7 targets. In order to evaluate this effect on physiological mRNA targets, we conducted RNAseq in WT, *Trim71* KO, *Lin28a* KO and double KO ESCs (Fig. 3A). Principal component analysis (PCA) showed four distinguishable clusters corresponding to each cell line, with *Lin28a* KO and double KO ESCs clustering close to each other (Fig. 3B-C). We then conducted co-expression network analysis (Fig. 3D) and found two clusters of genes which were consistently downregulated in all KO cell lines compared to WT ESCs (modules C and E, marked with a yellow asterisk in Fig. 3D). To evaluate the possibility that some of those downregulated genes had been repressed by let-7, we conducted an unbiased miRNA enrichment analysis *in silico* employing the online software ShinyGO v0.61^42^. This software identifies miRNA targets present in a list of genes and returns the top 100 miRNAs whose targets are significantly enriched in that list. Using the 1812 genes of modules C and E as input, we found the let-7 family among the top 100 enriched miRNA families (P-value (FDR) = 2.7E-07), with a total of 118 let-7 targets (≈ 6.5 %) found within modules C and E (Suppl. Table 1A). However, a limitation of this analysis is that ShinyGO takes into account only data from a single chosen database. In order to increase the robustness of our analysis, we then integrated data from eight different miRNA databases (see details in Methods) and found a total of 329 (≈ 18.2 %) predicted targets for the different let-7 family members present in the aforementioned modules (Fig. 3E and Suppl. Table 1B). GO enrichment analysis revealed that these let-7 targets mostly participate in mitotic-related functions (Supp. Fig. 5A-B). Collectively, our analysis indicated that a substantial part of the changes observed in the downregulated transcriptomes of each KO ESC line derive from the common upregulation of let-7 miRNAs, which results in the repression of multiple let-7 physiological mRNA targets with crucial functions for ESCs.

**Figure 3.**
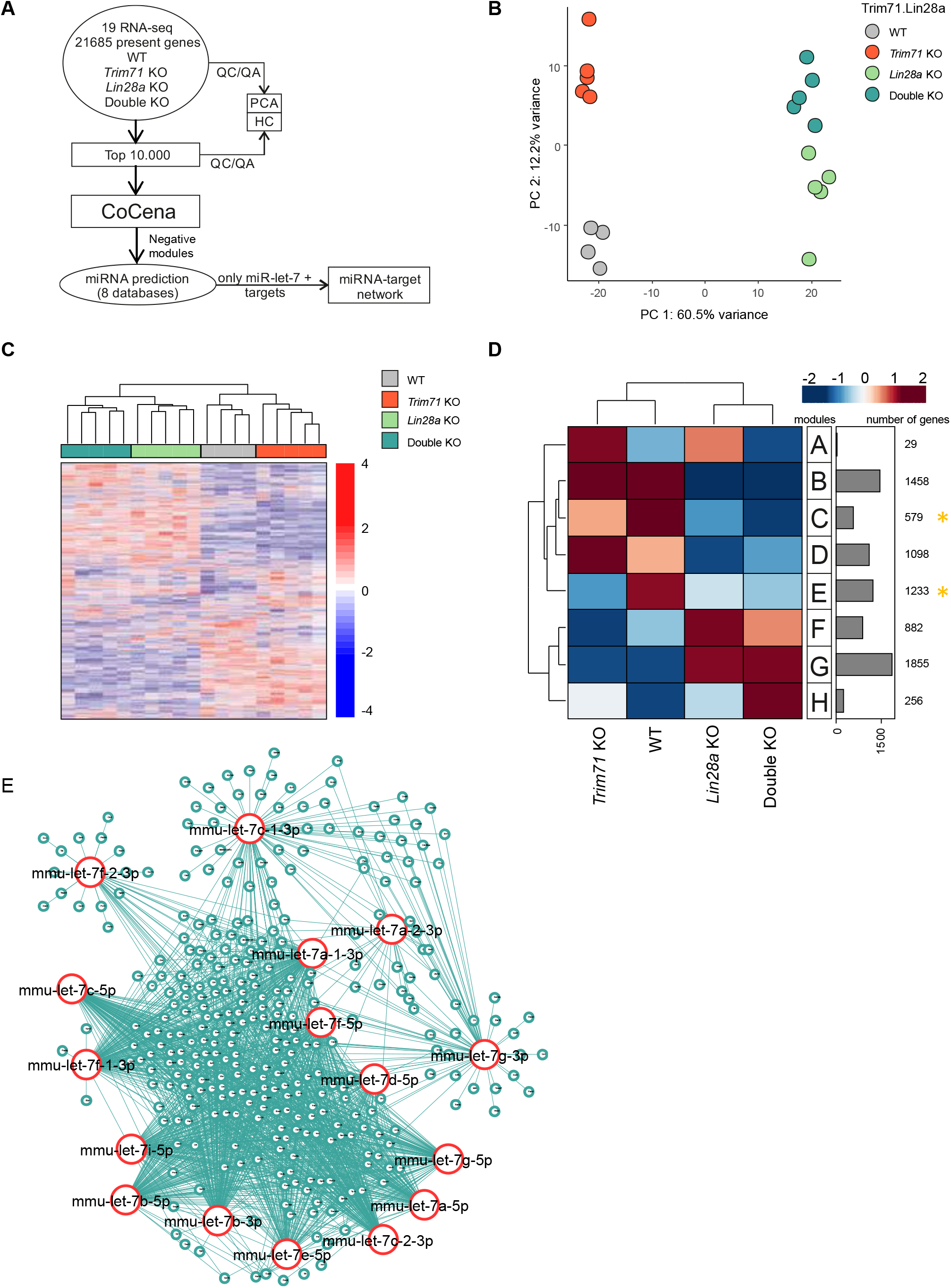
*Trim71* KO, *Lin28a* KO and double KO ESCs share the downregulation of multiple let-7 targets in their transcriptomic profiles. **A)** Schematic representation of the workflow for the transcriptomic analysis of ESCs. **B)** Principal component analysis (PCA) showing individual sequencing experiments for each genotype (n=4-5). **C)** Hierarchical clustering of the 10.000 most variable genes among the different ESC lines. **D)** Co-expression network analysis for the transcriptomes of the different ESC lines. Modules with commonly downregulated genes among all KO ESC lines are marked with a yellow asterisk and were used for E. **E)** miRNA network showing targets for the different let-7 family members found in modules C and E (marked with a yellow asterisk in D). See also Suppl. Fig. 5 and Suppl. Table 1.

### TRIM71 represses let-7 maturation through its interaction with the TUT4/LIN28 complex

After having found that TRIM71 cooperates with LIN28A in the repression of let-7 expression, we next aimed at elucidating the underlying molecular mechanism. A previous study reported an interaction between TRIM71 and the paralog protein LIN28B in HEK293T cells^27^. It was suggested that TRIM71 mediates the ubiquitylation and proteasomal degradation of LIN28B, thereby inducing let-7 upregulation^27^. We found that both wild type TRIM71 and the RING ubiquitylation mutant C12LC15A^14,27^ co-precipitated with endogenous LIN28B in HEK293T cells (Fig. 4A-B). Both TRIM71 variants also interacted with ectopically expressed LIN28A (Fig. 4C). However, we observed that overexpression of TRIM71 in HEK293T cells did not affect neither endogenous LIN28B protein nor mRNA levels (Fig. 4D-E), and instead resulted in the specific downregulation of let-7a/g-5p miRNAs – but not of the housekeeping miR-16 – (Fig. 4F), similar to our observations in ESCs. Thus, our results showed that TRIM71 does not promote LIN28B degradation, and instead suggested that TRIM71 can functionally cooperate with both LIN28 isoforms to repress the expression of let-7 miRNAs.

**Figure 4.**
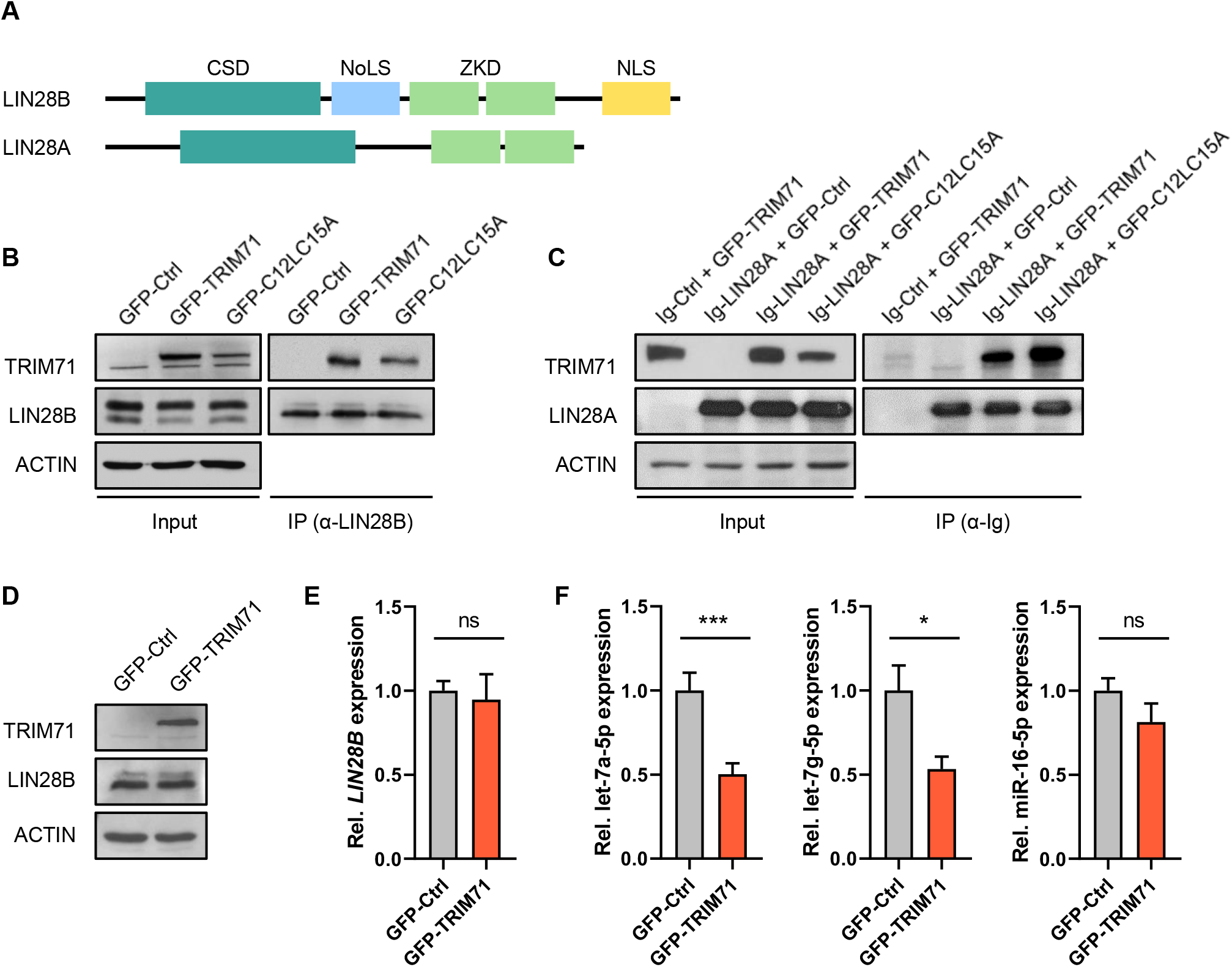
TRIM71 interacts with LIN28 proteins and specifically regulate let-7 miRNAs in HEK293T cells. **A)** Schematic representation of both LIN28 isoforms (LIN28A and LIN28B) domain organization. CSD = Cold Shock Domain; ZKD = Zinc Knuckle domain; NoLS = putative nucleolar localization sequence; NLS = Nuclear localization signal. **B)** Representative immunoblot showing GFP-tagged TRIM71 and C12LC15A co-precipitation with endogenous LIN28B in HEK293T cells. **C)** Representative immunoblot showing GFP-tagged TRIM71 and C12LC15A co-precipitation with Ig-tagged LIN28A overexpressed in HEK293T cells. Overexpression of an Ig-empty control vector together with wild type GFP-TRIM71 (first lane) excluded an Ig-mediated TRIM71-LIN28A interaction. Of note, GFP alone was not found co-precipitated with neither LIN28A nor LIN28B (data not shown), also excluding a possible GFP-mediated interaction. **D)** Representative immunoblot showing ectopic TRIM71 and endogenous LIN28B protein levels in HEK293 cells stably overexpressing GFP-Ctrl or GFP-TRIM71. **B)** RT-qPCR showing relative expression of LIN28B mRNA (n=3) and **C)** the indicated mature miRNAs (n=6) in HEK293 cells stably overexpressing GFP-Ctrl or GFP-TRIM71. RT-qPCR quantification of miRNAs and mRNAs was normalized to the levels of the housekeeping U6 snRNA and *HPRT1* mRNA, respectively. Error bars represent SD. ***P-value < 0.005, *P-value <0.05, ns = non-significant (unpaired student’s t-test). See also Suppl. Fig. 6.

Next, we designed several truncated constructs to map the interaction between TRIM71 and LIN28 proteins via co-precipitation experiments in HEK293T cells. We found that the Cold-shock domain (CSD) of LIN28 proteins (Suppl. Fig. 6A-B) and the NHL domain of TRIM71 (Suppl. Fig. 6C-E), respectively, are required and sufficient to establish their interaction in cell culture. We then produced a recombinant human TRIM71 NHL domain and evaluated its direct interaction with the LIN28A-pre-let7 complex *in vitro* via electrophoretic mobility shift assays (EMSA). We found LIN28A to specifically interact with pre-let-7 (K_D_ ≈ 55.5), but not with pre-miR-16 (Suppl. Fig. 7A-B). Strikingly, the addition of TRIM71’s NHL domain did neither result in a detectable shift of the LIN28A-pre-let-7 complex (PR) nor the pre-let-7 alone (R) at any of the tested NHL concentrations (Suppl. Fig. 7C-D). Based on these findings, we reasoned that other cellular components are required to mediate or stabilize the TRIM71-LIN28 interaction.

LIN28 proteins are known to induce pre-let-7 degradation through the recruitment of the terminal uridyl-transferase enzymes TUT4/7^33,40^. Thus, we first evaluated the ability of TRIM71 to interact with TUT4. Indeed, we found endogenous TUT4 co-precipitated with TRIM71 in control HEK293T cells, which lack LIN28A, as well as in LIN28B knockdown HEK293T cells (Fig. 5A). This indicated that LIN28 proteins are not indirectly mediating the interaction between TRIM71 and TUT4. In contrast, the binding between TRIM71 and endogenous LIN28B was abrogated upon TUT4 knockdown, suggesting that TUT4 facilitates the interaction between TRIM71 and LIN28B (Fig. 5B). The interaction between TRIM71 and TUT4/LIN28B was resistant to RNase treatment (Fig. 5C), demonstrating that these three partners form a stable protein complex. Consistent with these results, the NHL domain of TRIM71 was sufficient for TUT4 binding in HEK293T cells (Fig. 5D). Furthermore, the RING ubiquitylation mutant C12LC15A, which we had previously shown to also bind LIN28 proteins, was also found co-precipitated with TUT4 (Fig. 5E). Collectively, our data strongly support the formation of a tripartite TRIM71/TUT4/LIN28 complex which mediates the degradation of pre-let-7 miRNAs more efficiently than the TUT4/LIN28 complex alone.

**Figure 5.**
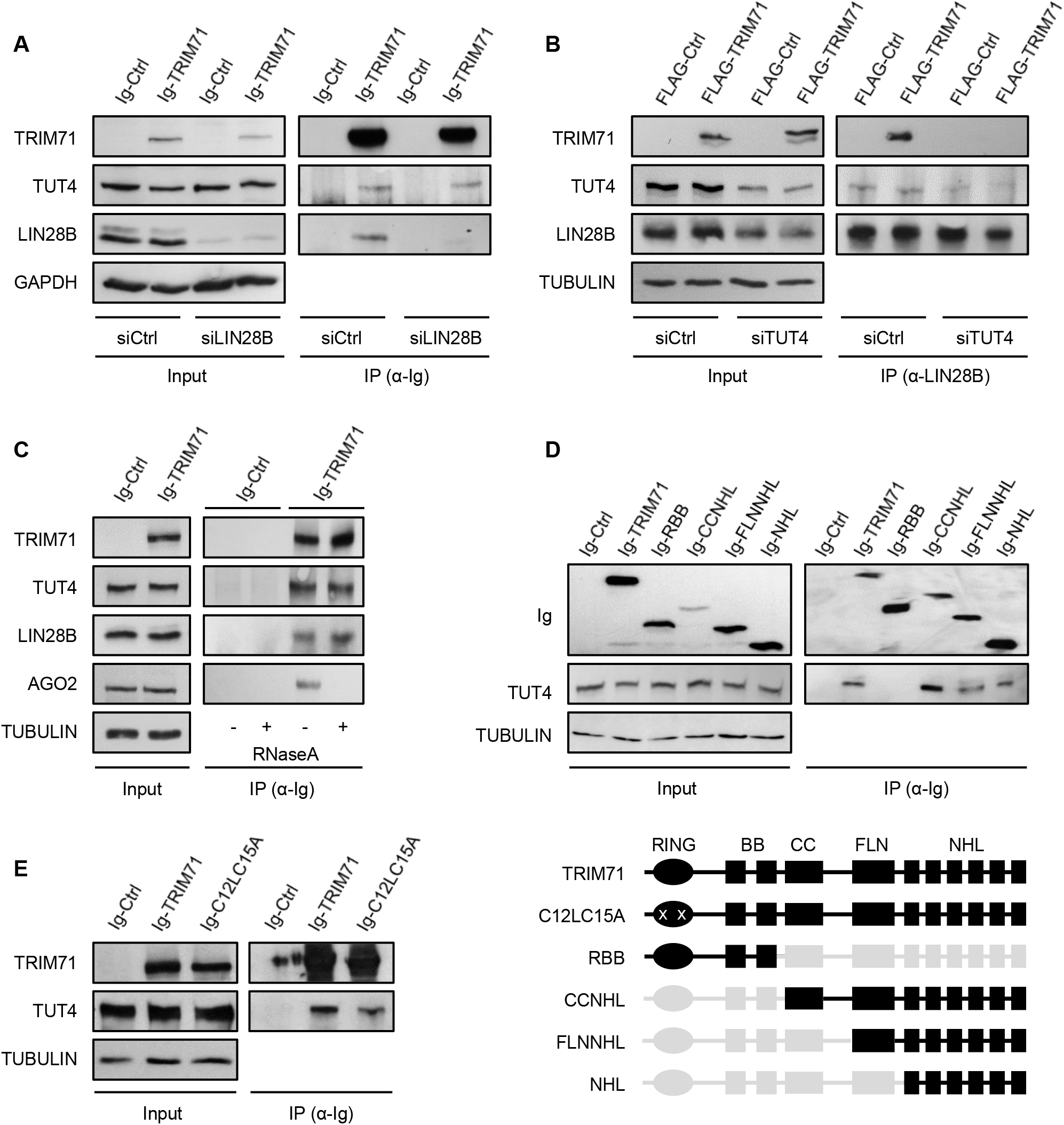
The interaction between TRIM71 and LIN28 is mediated by the uridylating enzyme TUT4. **A)** Representative immunoblot showing the co-precipitation of endogenous TUT4 with ectopically expressed Ig-TRIM71 in control HEK293T cells (siCtrl) – which lack LIN28A expression – and upon LIN28B knockdown (siLIN28B). **B)** Representative immunoblot showing the co-precipitation of ectopically expressed FLAG-TRIM71 with endogenous LIN28B in control (siCtrl) and TUT4 knockdown (siTUT4) HEK293T cells. **C)** Representative immunoblot showing the RNA-independent interaction between TRIM71 and TUT4 or LIN28B. AGO2 was used as a control of an RNA-dependent interaction. **D)** Representative immunoblot showing the co-precipitation of endogenous TUT4 with different Ig-tagged TRIM71 truncated proteins in HEK293T cells, depicted below. For each construct, present domains are depicted in black, deleted domains are depicted in grey and mutations are marked with a white “x”. **E)** Representative immunoblot showing the co-precipitation of endogenous TUT4 with Ig-tagged TRIM71 and C12LC15A overexpressed in HEK293T cells. See also Suppl. Fig. 7.

Since both TRIM71 and C12LC15A were able to interact with TUT4/LIN28, we then asked whether C12LC15A could also mediate let-7 downregulation. To this end, we evaluated the ability of wild type TRIM71 and RING ubiquitylation mutant C12LC15A to regulate let-7 expression and let-7 activity in several cell lines with different expression levels of LIN28 proteins: i) ESCs, which mostly express LIN28A, ii) HEK293T cells which exclusively express LIN28B, iii) the mouse embryonic fibroblast cell line NIH3T3 lacking expression of both LIN28 proteins, and iv) the human malignant T cell line Jurkat E6.1, in which both LIN28 proteins are lacking as well (Fig. 6A). Those cell lines expressing either LIN28A or LIN28B (ESCs and HEK293T, respectively) showed lower let-7a expression levels than cell lines lacking LIN28 proteins (NIH3T3 and Jurkat E6.1), as expected (Fig. 6B). Accordingly, let-7 activity was lower in LIN28-expressing cells, as shown by a higher de-repression of the let-7 luciferase reporter (Fig. 6C). Overexpression of both TRIM71 and C12LC15A resulted in a 60% downregulation of let-7 expression in ESCs, and a 25-30% downregulation of let-7 expression in HEK293T cells, proving that the RING domain (E3 ubiquitin ligase activity) of TRIM71 is not required for this function (Fig. 6D). In contrast, the overexpression of TRIM71 or C12LC15A in NIH3T3 and Jurkat E6.1 cells did not affect let-7 levels, which were only downregulated upon LIN28A overexpression (Fig. D). So far, our results have shown that TRIM71 establishes an RNA-independent interaction with the TUT4/LIN28 complex via its NHL domain, and enhances the repression of pre-let-7 maturation in an ubiquitylation-independent manner.

**Figure 6.**
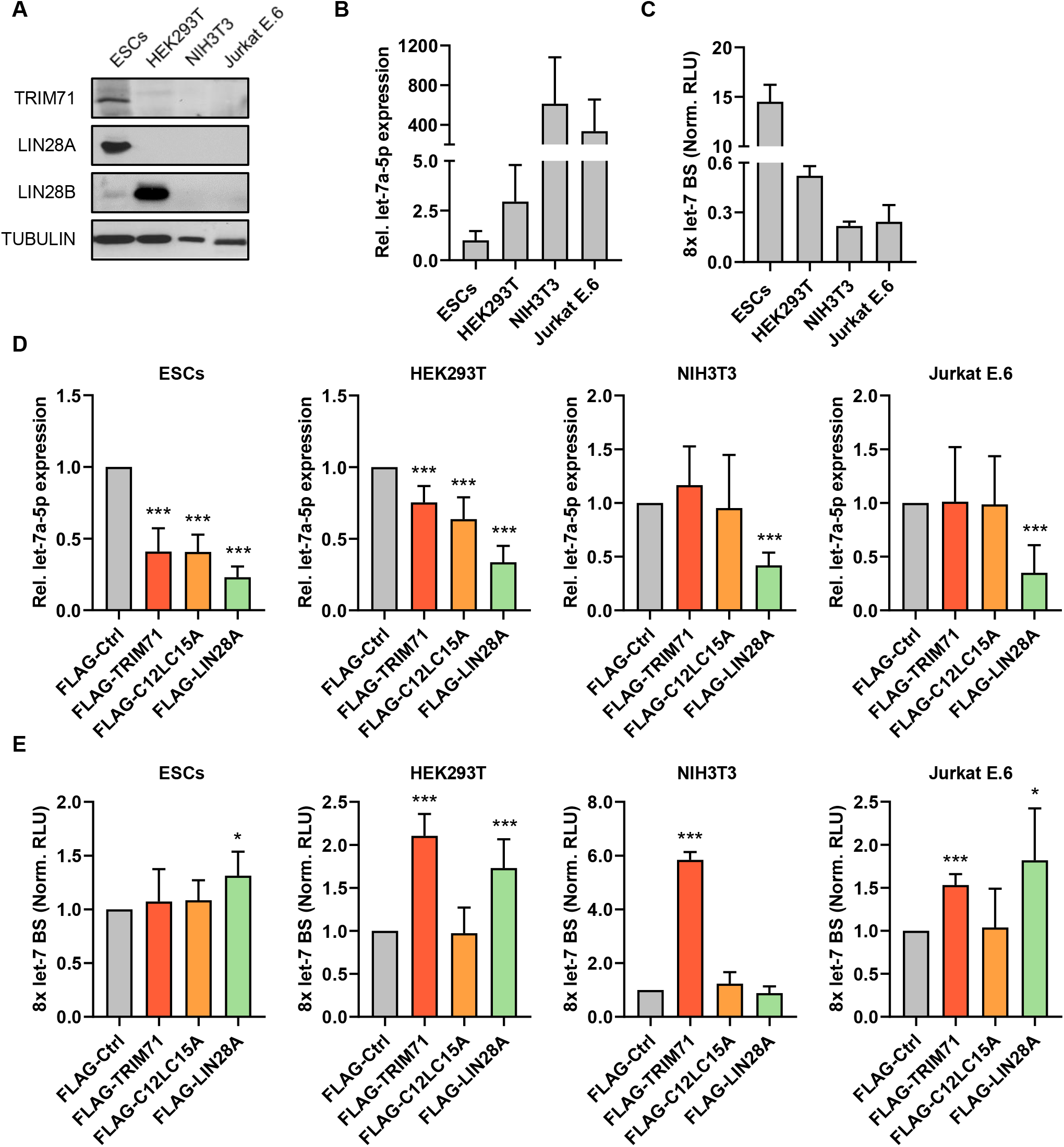
TRIM71 represses let-7 expression and activity via two independent mechanisms. **A)** Representative immunoblot showing the levels of endogenously expressed TRIM71 and LIN28 proteins in the indicated cell lines, used for experiments depicted in B-E. **B)** RT-qPCR showing the expression of mature let-7a miRNA in the indicated cell lines (n=3 6). **C)** Let-7 reporter assay upon transient transfection of a Renilla Luciferase reporter under the control of a 3’UTR containing 8x Let-7 binding sites (BS) in the indicated cell lines (n=3). Norm. RLU = Normalized Relative Light Units. **D)** RT-qPCR showing the expression of mature let-7a miRNA in the indicated cell lines upon overexpression of FLAG-tagged TRIM71, C12L15A and LIN28A (n=3-8). **E)** Let-7 reporter assays in the indicated cell lines upon overexpression of FLAG-tagged TRIM71, C12L15A and LIN28A (n=3-8). Norm. RLU = Normalized Relative Light Units. RT-qPCR quantification of let-7a was normalized to the levels of the housekeeping U6 snRNA. Error bars represent SD. ***P-value < 0.005, *P-value <0.05 (unpaired student’s t-test between FLAG-Ctrl and each other condition).

### TRIM71 regulates let-7 activity via a TUT4/LIN28-independent mechanism

When we analyzed let-7 activity in ESCs, HEK293T, NIH3T3 and JurkatE.6 cell lines via luciferase reporter assays, we observed striking discrepancies with the parallel let-7 expression measurements. Whereas both wild type TRIM71 and RING ubiquitylation mutant C12LC15A were able to downregulate let-7 expression to the same extent (Fig. 6D), only wild type TRIM71 induced a significant de-repression of the let-7 reporter (Fig. 6E) – from now on referred as repression of let-7 activity –. Furthermore, although a TRIM71-mediated downregulation of let-7 expression was only observed in LIN28-expressing cells (Fig. 6D), a TRIM71-dependent repression of let-7 activity was also found in LIN28-lacking cells (Fig. 6E). These results strongly suggested that TRIM71 employs distinct independent mechanisms for the regulation of let-7 expression and activity.

In line with this hypothesis, TRIM71 continued to repress let-7 activity after knockdown of either LIN28B or TUT4 in HEK293T cells (Fig. 7A-B and Suppl. Fig. 8A-B). Interestingly, TRIM71 was able to repress let-7 activity even upon overexpression of a mature let-7a miRNA duplex (Fig. 7A), indicating that this activity regulation mechanism occurs downstream of the expression regulation mechanism. To confirm this, we evaluated the repression of let-7 activity by TRIM71 and LIN28A upon overexpression of a pre-let-7a miRNA stem loop and a mature let-7a miRNA duplex (Fig. 7C). Both TRIM71 and LIN28 were able to relieve the repression of the let-7 reporter to a similar extent upon pre-let-7 overexpression. However, only TRIM71 was able to significantly relieve the repression of the let-7 reporter upon mature let-7a overexpression (Fig. 7C). These data showed that TRIM71-mediated let-7 activity regulation acts downstream of the TUT4/LIN28-dependent let-7 expression regulation, which operates at the precursor miRNA stage.

**Figure 7.**
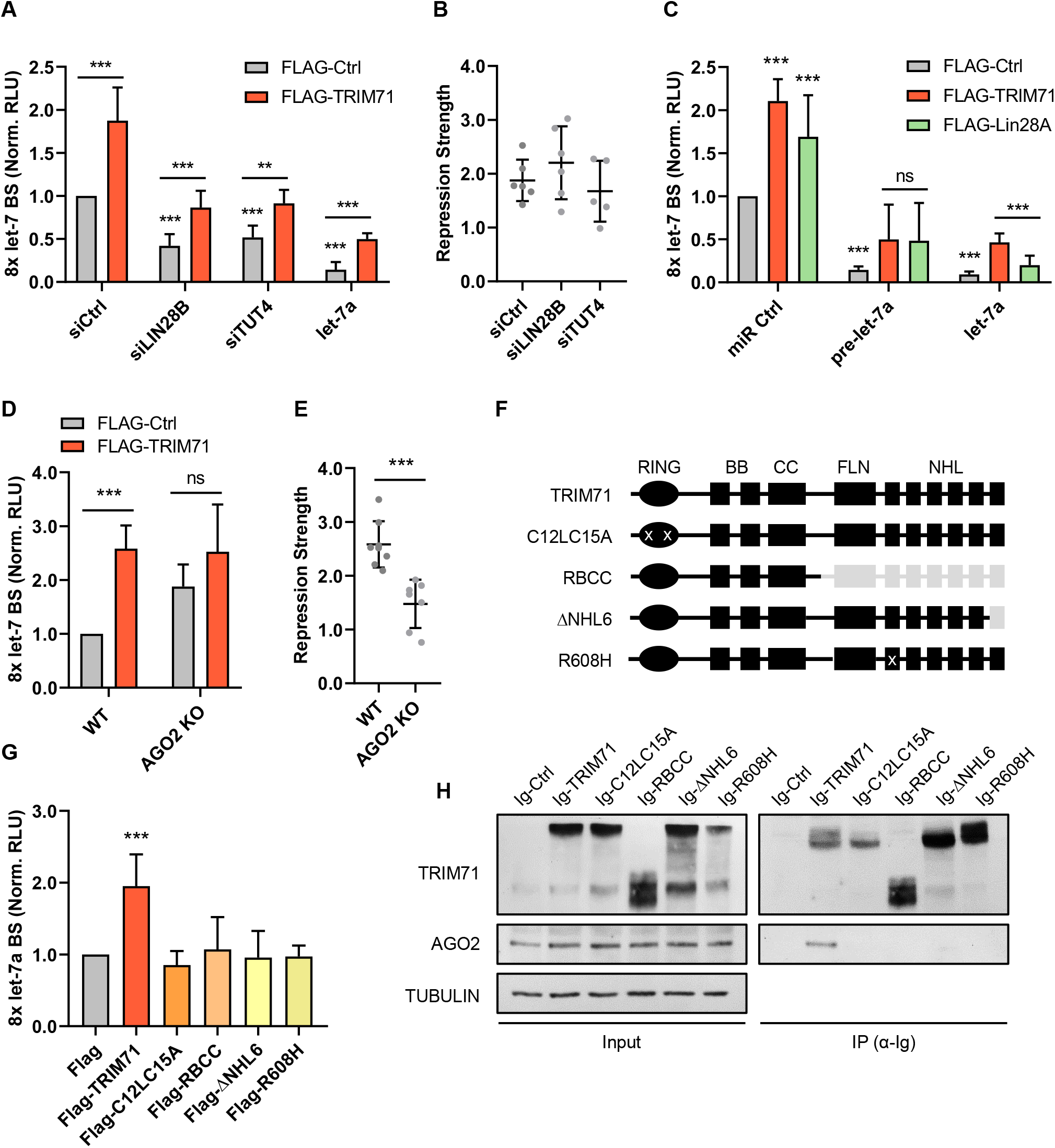
TRIM71 relies on AGO2 binding for the specific repression of Let-7 activity. **A)** Let-7 reporter assay upon overexpression of FLAG-Ctrl or FLAG-TRIM71 in control (siCtrl), LIN28B knockdown (siLIN28B) and TUT4 knockdown (siTUT4) HEK293T cells. Ectopic let-7 miRNA duplex expression was used as a control for efficient let-7 reporter repression, but revealed a regulation of mature let-7 activity (n=4-6). **B)** TRIM71-mediated let-7 activity repression strength, calculated from values in A, as Norm. RLU (FLAG-TRIM71) / Norm. RLU (FLAG-Ctrl) for each condition. **C)** Let-7 reporter assay upon FLAG-Ctrl, FLAG-TRIM71 or FLAG-LIN28A overexpression in HEK293T cells after ectopic expression of a control miRNA duplex (miR Ctrl) – representing the regulation of endogenously expressed let-7 –, a pre-let-7a stem loop or a mature let-7a miRNA duplex (n=3-8). **D)** Let-7 reporter assay upon overexpression of FLAG-Ctrl or FLAG-TRIM71 in wild type (WT) and AGO2 knockout (KO) HEK293T cells (n=6-8). **E)** TRIM71-mediated let-7 activity repression strength, calculated from values in D, as Norm. RLU (FLAG-TRIM71) / Norm. RLU (FLAG-Ctrl) for each condition. **F)** Schematic representation of TRIM71 constructs used in G and H. For each construct, present domains are depicted in black, deleted domains are depicted in grey and mutations are marked with a white “x”. **G)** Let-7 reporter assay in HEK293T cells upon overexpression of different FLAG-tagged TRIM71 constructs depicted in F (n=3-10). **H)** Representative immunoblot showing the co-precipitation of endogenous AGO2 with different Ig-tagged TRIM71 constructs – depicted in F – overexpressed in HEK293T cells. Norm. RLU = Normalized Renilla Light Units. Error bars represent SD. ***P-value < 0.005, **P-value <0.01, ns = non-significant (unpaired student’s t-test between the main control condition and each other condition, unless indicated by a line joining the two compared conditions). See also Suppl. Fig. 8.

Thus, our results show that TRIM71 regulates let-7 expression and activity via two independent, mechanistically discernable functions: on the one hand, TRIM71 relies on its interaction with the TUT4/LIN28/pre-let 7 complex to repress miRNA maturation, and thus downregulates let-7 expression; on the other hand, TRIM71 relies on its RING domain – E3 ubiquitin ligase function – to repress the activity of the mature let-7 miRNA in a TUT4/LIN28-independent manner.

### TRIM71 relies on AGO2 binding for the specific repression of let-7 activity

We next aimed to investigate how TRIM71 could repress let-7 miRNA activity. The activity of mature miRNAs is assisted by the RNA-Induced Silencing Complex (RISC). Argonaute (AGO) proteins - particularly the well-characterized AGO2 – are the major effectors of the RISC, since they bind the miRNA duplex, identify the guide strand, and find the complementary mRNA to repress its translation and often induce its degradation^43^. Several TRIM-NHL proteins, including TRIM71, are known to interact with AGO proteins in various species^11,14,20,21,24,28,44^. To evaluate whether TRIM71-mediated let-7 activity repression is linked to AGO2 function, we conducted let-7 reporter assays in wild type (WT, AGO^+/+^) and AGO2 knockout (KO, AGO^−/−^) (Suppl. Fig. 8C)^24^ HEK293T cells. Indeed, we found a diminished TRIM71-mediated repression of let-7 activity in AGO2 knockout HEK293T cells (Fig. 7D-E). Although this effect could be attributable to a general diminished miRNA activity in AGO2 KO cells, these cells continued to downregulate the let-7 target *HMGA2* upon overexpression of mature let-7 (Suppl. Fig. 8D), suggesting that other AGO proteins may take over miRNA-mediated silencing in the absence of AGO2 as it was previously described^45^.

The interaction between TRIM71 and AGO2 has been mapped to the NHL domain and is known to be RNA-dependent^20,21,24^ (as we have also shown in Fig. 5C). We therefore evaluated the ability of the of several TRIM71 NHL domain mutants with impaired RNA binding ability (RBCC^21^, ΔNHL6^24^ and R608H^12,23^), as well as of the RING ubiquitylation mutant C12LC15A, to bind AGO2 and to repress let-7 activity. We found that all investigated TRIM71 mutants failed to bind AGO2 and regulate let-7 activity (Fig. 7F-H). This correlation strongly suggested that TRIM71 depends on AGO2 for the inhibition of let-7 miRNA activity.

A previous study postulated that TRIM71 inhibits miRNA activity through a reduction of AGO2 stability via ubiquitylation^14^. However, later publications have repeatedly shown in various systems that TRIM71 does not induce changes in AGO2 stability^11,16,20,21,24^ (see also Fig. 5C and 7H). Accordingly, luciferase reporter assays for other miRNAs showed that TRIM71 does not repress miRNA activity globally, as the activity of miR-16 or miRNA-19 remained unaffected upon TRIM71 overexpression in HEK293T cells (Suppl. Fig. 8E). Collectively, our data indicate that TRIM71 represses the activity of let-7 miRNAs in an AGO2-dependent manner. To interact with AGO2, TRIM71 requires both an intact NHL domain and a functional RING domain. Such an interaction neither results in decreased AGO2 stability nor in a global repression of miRNA activity.

### TRIM71 binds and stabilizes let-7 mRNA targets in ESCs

TRIM71 is known to interact and/or co-localize with several RISC-associated factors, namely DICER^14^, MOV10 and TNRC6B^21^. Furthermore, TRIM71 is an mRNA-binding protein^21,23,24^, and we have shown the interaction between TRIM71 and AGO2 to be RNA-dependent. Thus, we hypothesized that the miRNA-inhibitory function of TRIM71 might take place on active RISCs, and that TRIM71 interacts with AGO2/RISC via its binding to specific mRNAs. To test this hypothesis, we asked whether TRIM71 binds and stabilizes let-7 targets in ESCs. To this end, we analyzed a published data set which includes RNA-seq data from TRIM71 RNA-IP experiments in ESCs, accompanied by transcriptomics on several *Trim71* mutant ESCs, including *Trim71* KO ESCs, *Trim71* RING mutant ESCs and *Trim71* NHL mutant ESCs^23^. RING mutant ESCs harbor the E3 ligase-impairing mutation C12LC15A^14,27^, and NHL mutant ESCs contain an R738A point mutation which was previously shown to impair TRIM71’s ability to bind RNA^46^.

Since we have shown that both the RING and the NHL domains are required for AGO2 binding and let-7 activity repression, we expected let-7 targets regulated by TRIM71 via the AGO2-dependent mechanism to be commonly downregulated in all mutant ESCs. Co-expression network analysis of transcripts derived from WT, *Trim71* KO, RING mutant, and NHL mutant ESCs showed four clusters of genes commonly downregulated in all mutant ESCs (modules B, C, J and K, marked with a yellow asterisk) (Fig. 8A-B). We then used the 2397 genes corresponding to all four modules for miRNA enrichment analysis via ShinyGO and found the let-7 family to be among the top 100 enriched miRNA families (P-value (FDR) = 2.54E-04), with a total of 131 let-7 targets (≈ 5.5 %) identified within modules B, C, J and K (Suppl. Table 2A). Integrating information from eight different databases increased this number to 359 predicted let-7 targets (≈ 15 %) (Fig. 8A, Fig. 8D and Suppl. Table 2B). These results recapitulated with a new dataset our previous findings showing that TRIM71 depletion results in the downregulation of multiple let-7 targets in ESCs.

**Figure 8.**
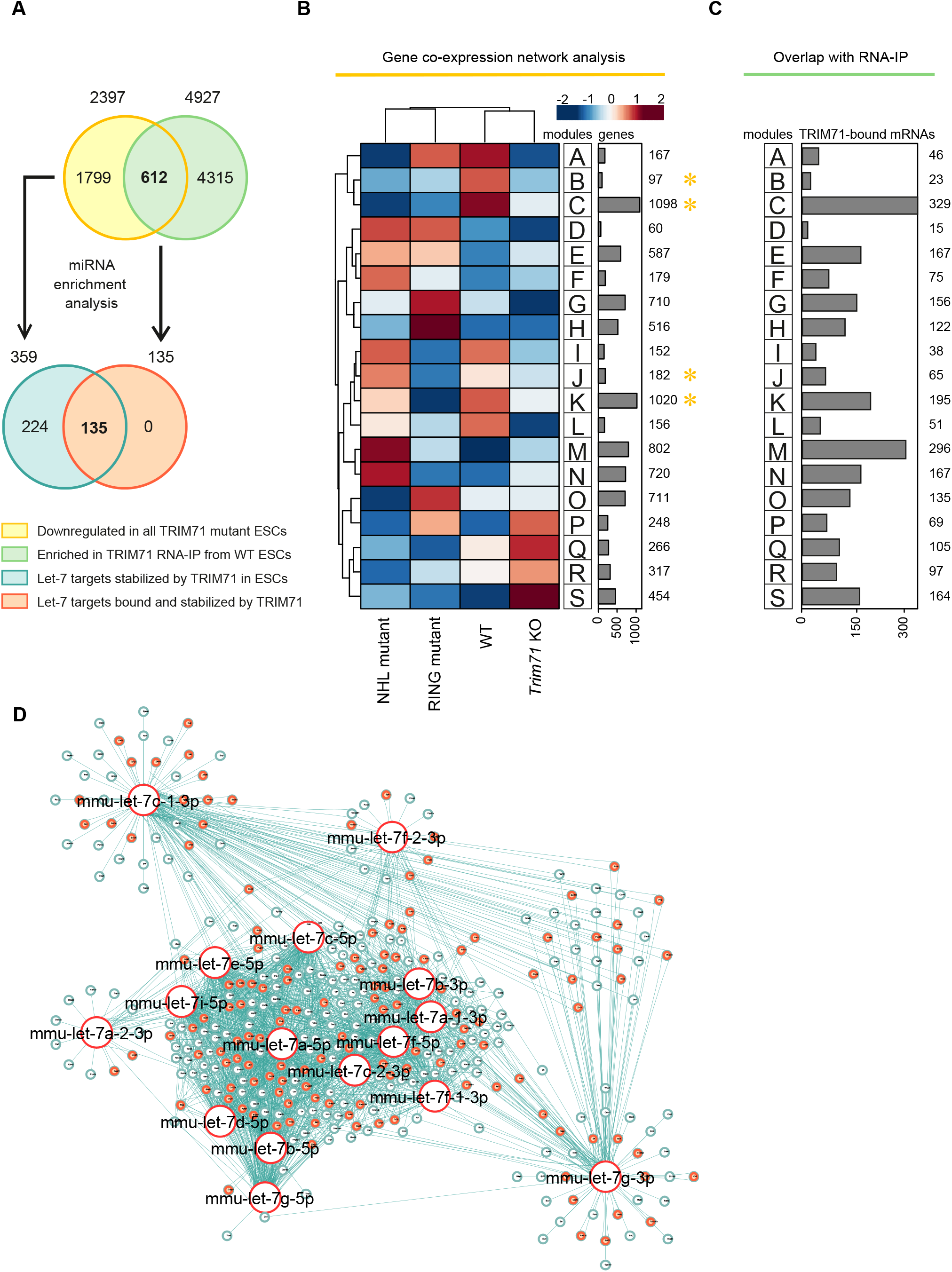
TRIM71 directly binds and stabilizes let-7 targets in ESCs. **A)** Schematic representation for the analysis conducted in B-D. Genes commonly downregulated in all *Trim71* mutant ESCs (yellow) – obtained from B – are enriched for let-7 targets (blue) (Suppl. Table 2). Genes found enriched (>2-fold) in TRIM71 RNA-IPs in ESCs (green) were overlapped with genes commonly downregulated in all *Trim71* mutant ESCs (yellow) – overlap depicted in C – to find genes bound and stabilized by TRIM71 in ESCs. For those genes, miRNA enrichment analysis was conducted (Suppl. Table 3) to identify let-7 targets bound and stabilized by TRIM71 (orange). **B)** Co-expression network analysis for the indicated ESC lines using an available dataset (GSE134125)^23^ revealed genes commonly downregulated in all *Trim71* mutant ESCs (modules B, C, J and K, marked with a yellow asterisk). **C)** Overlap of TRIM71 RNA-IP data in ESCs (GSE134125)^23^ with gene expression data, showing the number of TRIM71-bound mRNAs in ESCs within each module from B. **D)** miRNA network showing targets for the different let-7 family members found among commonly downregulated genes in all *Trim71* mutant ESCs (modules B, C, J and K from B), with let-7 targets directly bound by TRIM71 in ESCs marked in orange. See also Suppl. Tables 2 and 3.

We then overlapped the gene expression data with the RNA-IP data to find TRIM71-bound mRNAs within the aforementioned four clusters of genes, i.e., mRNAs bound and stabilized by TRIM71 (Fig. 8A-C). This approach identified 612 mRNAs bound and stabilized by TRIM71. Using these mRNAs for miRNA enrichment analysis via ShinyGO, we detected again the let-7 family among the top 100 enriched miRNA families (P-value (FDR) = 1.20E-05), with a total of 48 let-7 targets (≈ 7.8 %) within all mRNAs bound and stabilized by TRIM71 (Suppl. Table 3A). Integrating information from eight different databases increased this number to 135 predicted let-7 targets (≈ 22 %) (Fig. 8A, Fig. 8D and Suppl. Table 3B). This reveals that 37.6 % (135) of all let-7 targets stabilized in the presence of TRIM71 (359) are directly bound by TRIM71.

Altogether, our analysis shows that TRIM71 interacts with specific let-7 mRNA targets, resulting in their stabilization, and that both the RING and the NHL domains of TRIM71 are required for the regulation of these let-7 targets. Having previously shown that both the RING and the NHL domains are required for AGO2 binding and let-7 activity regulation, our results strongly suggest that TRIM71 represses let-7 activity by inhibiting AGO2 function on active RISCs, to which TRIM71 associates via the interaction with specific mRNAs, i.e., let-7 targets.

### TRIM71 binds and stabilizes let-7 mRNA targets in HCC cells

TRIM71 is upregulated in patients with hepatocellular carcinoma (HCC) and its expression is correlated with advanced tumor stages and poor prognosis^24,44^. Several studies have shown that TRIM71 promotes proliferation in several HCC cell lines, including HepG2, Hep3B and Huh7^24,44,47^. In one of these studies, an increase of let-7 activity was observed in TRIM71 knockdown cells^44^. This effect was attributed to the previously reported global inhibition of miRNA activity caused by TRIM71-mediated ubiquitylation and proteasomal degradation of AGO2^14^. However, we did not find any evidence supporting a role for TRIM71 in the regulation of AGO2 stability.

Thus, we first evaluated AGO2-TRIM71 interaction in wild type HepG2 cells (Suppl. Fig. 8F), as well as AGO2 stability (Suppl. Fig. 8G) and let-7 activity (Suppl. Fig. 8H) upon TRIM71 knockdown in HepG2 cells. Our results showed that TRIM71 binds AGO2 and regulates let-7 activity in HepG2 cells without affecting AGO2 protein stability (Suppl. Fig. 8F-H). We then used an available transcriptomic dataset of *TRIM71* KO Huh7 cells^23^, and conducted co-expression network analysis (Fig. 9A-B). We then selected significantly downregulated mRNAs in *TRIM71* KO cells (2903 genes from modules A-E, marked with a yellow asterisk in Fig. 9B), and conducted miRNA enrichment analysis via ShinyGO. Again, we found the let-7 family to be among the top 100 enriched miRNA families (P-value (FDR) = 9.89E-04), with a total of 216 downregulated let-7 targets (≈ 7.4 %) (Suppl. Table 4A). Integrating information from eight different miRNA data bases increased this number to 282 predicted targets (≈ 9.7 %) for the members of the let-7 family (Fig. 9A, Fig. 9D and Suppl. Table 4B).

**Figure 9.**
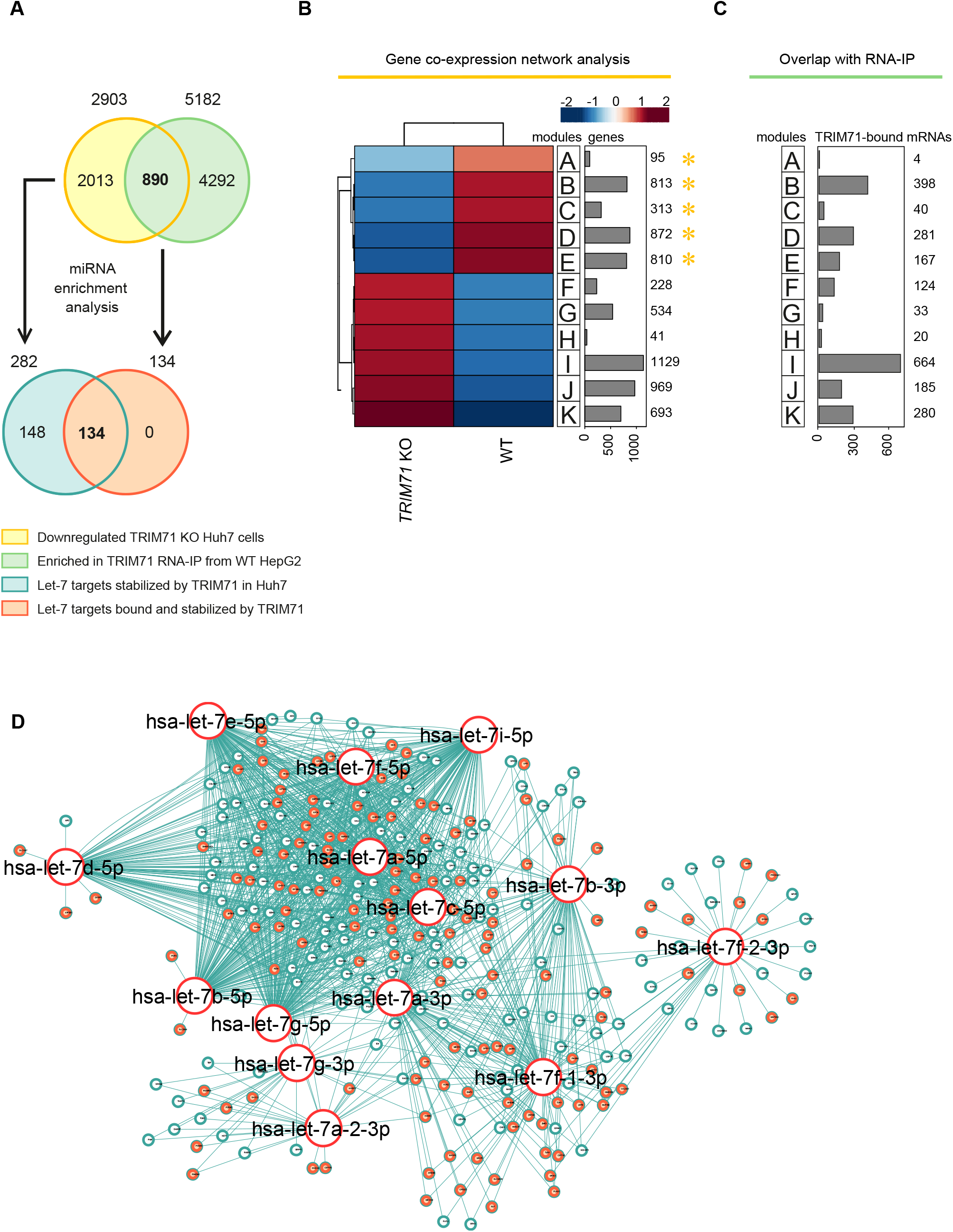
TRIM71 directly binds and stabilizes let-7 targets in HCC cells. **A)** Schematic representation for the analysis conducted in B-D. Genes downregulated in *TRIM71* KO Huh7 cells (yellow) – obtained from B – are enriched for let-7 targets (blue) (Suppl. Table 4). Genes found enriched (>2-fold) in TRIM71 RNA-IPs in HepG2 cells (green) were overlapped with genes downregulated in *TRIM71* KO Huh7 cells (yellow) – overlap depicted in C – to find genes bound and stabilized by TRIM71. For those genes, miRNA enrichment analysis was conducted (Suppl. Table 5) to identify let-7 targets bound and stabilized by TRIM71 in HCC cells (orange). **B)** Co-expression network analysis for WT and *TRIM71* KO Huh7 cells using an available dataset (GSE134125)^23^ revealed genes downregulated in *TRIM71* KO Huh7 cells (modules A-E marked with a yellow asterisk). **C)** Overlap of TRIM71 RNA-IP data in HepG2 cells^47^ with gene expression data, showing the number of TRIM71-bound mRNAs in HCC cells within each module from B. **D)** miRNA network showing targets for the different let-7 family members found among commonly downregulated genes in in *TRIM71* KO Huh7 cells (modules A-E), with let-7 targets directly bound by TRIM71 in HCC cells marked in orange. See also Suppl. Tables 4 and 5.

To evaluate whether TRIM71 was also able to interact with some of these let-7 targets in hepatocellular carcinoma cells, we used another available TRIM71 RNA-IP dataset from HepG2 cells^47^, and overlapped it with genes found to be downregulated in *TRIM71* KO Huh7 cells (Fig. 9A-C). We found a total of 890 genes bound and stabilized by TRIM71 in HCC cells, 87 of which (≈ 9.8 %) were found to be let-7 targets via miRNA enrichment analysis (P-value (FDR) = 1.30E-05) (Suppl. Table 5A). Integrating information from eight different miRNA data bases increased this number to a total of 134 predicted targets (≈ 15 %) for the different let-7 family members (Fig. 9A, Fig. 9D and Suppl. Table 5B). This shows that 47.5 % (134) of all let-7 targets stabilized in presence of TRIM71 (282) in HCC cells are also directly bound by TRIM71. While we propose that these targets are regulated by TRIM71 via the AGO2-dependent mechanism, let-7 targets stabilized but not bound by TRIM71 are susceptible to be regulated via the TUT4/LIN28-dependent mechanism, both in ESCs and HCC cells. Altogether, these results indicate that the TRIM71-mediated let-7 regulatory mechanisms reported by our work are not restricted to ESCs but also found to be operative in hepatocellular carcinoma cells.

## Discussion

Regulation of miRNAs is one of the major modulating mechanisms to fine-tune protein expression and subsequently protein function. TRIM-NHL proteins have been linked to the regulation of the miRNA pathway in several species^28^. In the present work we performed a comprehensive analysis of TRIM71-dependent regulation of miRNA expression and activity, and specifically focused on the regulation of the let-7 miRNA family.

By analyzing the miRNome of TRIM71-deficient murine ESCs^11^, we identified let-7 family members as some of the strongest upregulated miRNAs as compared to wild type ESCs. Mature miRNA duplexes of several let-7 members were found upregulated in *Trim71* KO ESCs, whereas expression of pri- and pre-let-7 were unaltered, suggesting that TRIM71 interferes with pre-let-7 maturation. Since LIN28 proteins are known to assist pre-let-7 degradation^32,33^, we generated *Lin28a* KO and *Trim71-Lin28a* double KO cells, and found an upregulation of mature let-7 miRNAs – but again not of pri- and pre-miRNA molecules – in all KO cell lines. Accordingly, all KO cell lines shared a downregulation of multiple let-7 targets in their transcriptomic profiles, most of which participate in mitotic functions.

Importantly, the *Trim71-Lin28a* double KO ESC line did not show any additional let-7 upregulation to that observed in single *Lin28a* KO ESCs, suggesting that TRIM71 depends on LIN28A for the regulation of let-7 expression. Indeed, overexpression of TRIM71 resulted in a significant downregulation of let-7 only in WT, but not *Lin28a* KO ESCs, confirming TRIM71’s dependency on LIN28A for this function. We found that TRIM71 can likewise functionally cooperate with the paralog protein LIN28B for the repression of let-7 expression in HEK293T cells. We demonstrated that TRIM71 is able to interact with both LIN28 isoforms, and found the TRIM71-LIN28 interaction to be RNA-independent, E3 ligase-independent, and mediated by TRIM71’s NHL domain and LIN28 protein’s CSD, respectively. Further analysis revealed that the uridylating enzyme TUT4 facilitates the interaction between TRIM71 and LIN28 proteins. Accordingly, the interaction between TRIM71 and TUT4 was also RNA-independent, E3 ligase-independent and mediated by the TRIM71’s NHL domain. Altogether, we showed that TRIM71 represses let-7 expression in cooperation with the TUT4/LIN28 complex, a well-established inhibitor of pre-let-7 maturation^33^.

Interestingly, a similar effect on pre-let-7 regulation was previously described for TRIM25, another member of the TRIM protein family that lacks an NHL domain^48^. In this case, TRIM25 was found to directly interact via its coiled-coil domain with the pre-let-7 stem loop and to enhance TUT4-mediated pre-let-7 uridylation. The role of TRIM25 as an E3 ligase in this context was not investigated and the exact mechanism of uridylation enhancement remains to be elucidated^48^. Nevertheless, this mechanism revealed the existence of miRNA-specific cofactors that stimulate TUT4-mediated uridylation. Thus, TRIM71 could also enhance pre-let-degradation by stimulating TUT4-mediated uridylation.

Our EMSA experiments showed that, unlike TRIM25^48^, TRIM71 is not able to directly interact with pre-let-7 miRNAs. This result is in line with a previous study that investigated ESC-specific miRNA-binding proteins, and identified TRIM71 as one of them for its binding to pre-miR-1 and pre-miR-29a, but not pre-let-7 species^26^. Thus, an interaction between TRIM71 and pre-let-7 is only indirectly mediated by the TUT4/LIN28 complex. Therefore, TUT4 and LIN28, but not TRIM71, are responsible for pre-miRNA target specificity. Hence, we propose that TRIM71 may be able to enhance the degradation of other pre-miRNAs reported to be regulated by the TUT4/LIN28 axis. Indeed, TRIM71-deficient ESCs showed an upregulation of miR-9 and miR-200 species^11^, both of which are pro-differentiation and tumor suppressor miRNAs whose precursors are targeted for degradation via the TUT4/LIN28 complex^33,49,50^. Notably, both miR-9 and miR-200 families were also present among the top 100 enriched miRNAs in all of our miRNA enrichment analysis. Thus, our study also provides several miRNA candidates which might be regulated in the same fashion as let-7.

Further supporting a TRIM71/TUT4/LIN28-mediated let-7 regulation mechanism, the overexpression of TRIM71 resulted in decreased let-7 expression in ESCs and HEK293T cells, which express LIN28A and LIN28B, respectively, but not in NIH3T3 or Jurkat E6.1 cells, which lack LIN28A/B expression. Notably, TRIM71-mediated let-7 downregulation was stronger in ESCs than in HEK293T cells. This difference might be explained by the distinct LIN28 isoforms expressed in either of these cell lines. While both LIN28 isoforms can regulate let-7 in the cytoplasm in a TUT4-dependent manner^32,33,51^, LIN28B can additionally block let-7 biogenesis in a TUT4-independent manner within the nucleus, where it binds pri- and pre-let-7 molecules preventing further processing and cytoplasmic export^32,36^. Given the cytoplasmic localization of TRIM71^14,20,24^, the efficiency of TRIM71-mediated let-7 regulation in LIN28B-expressing cells may be limited by the amount of LIN28B available in the cytoplasm, which is known to vary among cell types^36,52,53^.

Importantly, we provided evidence that the E3 ligase function of TRIM71 is dispensable for the regulation of let-7 expression, as the RING ubiquitylation mutant C12LC15A could interact with TUT4/LIN28 and repress let-7 expression to a similar extent as the wild type TRIM71. In contrast, only wild type TRIM71, but not C12LC15A, was able to repress let-7 activity in luciferase reporter assays. Furthermore, TRIM71-mediated repression of let-7 activity was also observed in NIH3T3 and Jurkat E6.1 cells in which let-7 expression was unaltered. These results revealed that TRIM71 employs two independent molecular mechanisms for the regulation of let-7 expression and activity. Further analysis confirmed that the regulation of let-7 activity occurred independently and downstream of the TUT4/LIN28 function.

An earlier study had identified TRIM71 as a global inhibitor of miRNA activity, a role that TRIM71 supposedly achieves via AGO2 protein ubiquitylation and proteasomal degradation^14^. However, we and others^11,16,20,21,24^ did not detect TRIM71-mediated changes of AGO2 stability. Accordingly, we observed a miRNA-specific rather than a global inhibition of miRNA activity upon TRIM71 overexpression. Nevertheless, we found that TRIM71-mediated repression of let-7 activity was impaired in AGO2 knockout HEK293T cells. Furthermore, we observed an impaired ability to repress let-7 activity in all TRIM71 mutants which failed to interact with AGO2. These results strongly suggested that TRIM71-mediated let-7 activity repression is linked to AGO2 function.

Other TRIM-NHL proteins, namely drosophila orthologues Brat, Mei-P26 and Wech/Dappled, *C. elegans* NHL-2 and mammalian TRIM32, were all found to interact with AGO proteins, but in neither case such an interaction resulted in reduced AGO protein stability^21,54–56^. Interestingly, *C. elegans* NHL-2 and mammalian TRIM32 were also found to affect miRNA activity^55,56^. Specifically, both NHL-2 and TRIM32 were found to enhance the activity of let-7 miRNAs and to promote cell differentiation^55,56^. Although TRIM71- and TRIM32-mediated miRNA activity regulation result in opposite functional outcomes, their molecular mechanisms seem to be highly similar: both proteins interact with AGO proteins via their NHL domain^20,21,56^ and affect the activity of specific miRNAs, including let-7. Furthermore, both TRIM32 and TRIM71 were found to induce AGO2 ubiquitylation without leading to their degradation^21^, suggesting that ubiquitylation-dependent events may regulate AGO2 function without affecting its stability.

Several post-translational modifications are known to alter AGO2 function^57,58^. For instance, AGO2 phosphorylation was shown to reduce let-7 activity by causing the dissociation of let-7 from the RISC complex, resulting in the stabilization of let-7 targets^59^. Similarly, TRIM71-mediated AGO2 ubiquitylation could lead to allosteric inhibition of AGO2 binding to specific miRNAs, resulting in the repression of miRNA activity. Alternatively, TRIM71 autoubiquitylation could be required for AGO2 binding. This type of autoregulation has been observed for TRIM32, as autoubiquitylation is required for its localization into cytoplasmic processing bodies^60^. Supporting this second hypothesis, our results showed that the TRIM71 ubiquitylation mutant C12LC15A not only failed to repress let-7 activity, but also to interact with AGO2.

The interaction of TRIM71 with AGO2 required not only a functional RING domain, but also an intact NHL domain. We showed the interaction between TRIM71 and AGO2 to be RNA-sensitive, and TRIM71 NHL mutants with impaired RNA-binding capacity failed to bind AGO2. Thus, we hypothesized that TRIM71 binds AGO2 on active RISCs through its interaction with specific let-7 target mRNAs. This may also explain why TRIM71 does not affect global miRNA activity, as its miRNA specificity would be determined by the TRIM71-bound mRNA target. Indeed, we found that TRIM71 interacts with multiple let-7 targets in mouse ESCs and human HCC cells, resulting in their stabilization. Importantly, TRIM71 required both the RING and NHL intact domains for the stabilization of those targets, again suggesting that the TRIM71-AGO2 interaction is required for the repression of let-7 activity. Thus, our data collectively support a role for TRIM71 as an inhibitor of let7-associated RISCs.

A very recent study has shown that TRIM71 is able to bind and stabilize specific mRNAs in the HCC cell line HepG2^47^. Beyond the fact that we found some of these mRNAs to be let-7 targets, our work also confirms that TRIM71 is capable of positively regulating mRNA expression post-transcriptionally in ESCs. In this regard, further research should clarify which mechanisms govern the distinct fates – degradation or enhanced stability – of TRIM71-bound mRNAs.

To sum up, TRIM71 was first discovered more than 20 years ago as a mRNA target of the miRNA let-7. In the present study, we have uncovered a role for TRIM71 as a let-7 repressor, revealing a negative feedback regulation between these two highly conserved developmental cues. Our work shows that TRIM71 modulates the let-7 miRNA pathway via specific interactions with two distinct protein complexes, TUT4/LIN28 and AGO2/mRNAs (RISC), in order to regulate let-7 expression and let-7 activity, respectively. These mechanisms induced significant transcriptomic changes in ESCs and HCC cells, suggesting that TRIM71-mediated miRNA regulatory mechanisms play important roles both in developmental and oncogenic processes. Future studies may reveal further molecular requirements of the mechanisms described by our work, as well as shed further light on the precise *in vivo* implications of TRIM71-mediated miRNA regulation during embryogenesis and tumorigenesis.

## Methods

### Cell lines and cell culture

The generation of wild type mouse ESCs (WT, *Trim71^fl/fl^*) from conditional *Trim71* full knockout mouse (*Trim71^fl/fl^; Rosa26-CreERT2*) was described in our previous work^11^. These cells were then used to generate *Lin28a* KO ESCs (*Trim71^fl/fl^; Lin28a^−/−^*) via gene editing using transcription activator-like effectors nucleases (TALENs) as previously described^61^. The TALEs were engineered to bind specific DNA sequences in the *Lin28a* locus (5’-GGGGCCCGGGGCCACGGGC-3’ and 5’-GCTGGTTGGACACCGAGCC-3’) and were fused to the unspecific FokI endonuclease catalytic domain to generate a double strand break close downstream of the *Lin28a* start codon. A donor DNA with *Lin28a* homology arms was designed to insert a G-418 resistance gene within the *Lin28a* locus by homologous recombination in order to allow pre-selection of edited clones, while disrupting *Lin28a* coding sequence.

*Trim71* knockout ESCs (KO, *Trim71^−/−^*) and *Trim71-Lin28a* double KO ESCs (*Trim71^−/−^; Lin28a^−/−^*) were generated from WT (*Trim71^fl/fl^*) and *Lin28a* KO ESCs (*Trim71^fl/fl^; Lin28a^−/−^*), respectively, by addition of 500 nM of 4-hydroxytamoxifen (4-OHT) in their culture media for 48 h, followed by further culture for 72 h to achieve full protein depletion. The generation of HEK293 cells stably overexpressing GFP or GFP-TRIM71, and of AGO2 KO HEK293T cells was described in our previous work^24^.

ESCs were cultured in 0.1 % gelatin-coated dishes and maintained in 2i+LIF media (DMEM knockout media supplemented with 15 % FCS, 1 % Penicillin-Streptomycin, 0.1 mM NEAA, 2 mM L-GlutaMAX, 100 μM β-mercaptoethanol, 0.2 % in-house produced LIF (supernatant from L929 cells), 1 μM of MEK/ERK inhibitor PD0325091 and 3 μM of GSK-3 inhibitor CHIR99021). HEK293(T) cells and NIH3T3 cells were maintained in DMEM media supplemented with 10 % FBS and 600 μg/ml of G418 antibiotic solution (HEK293) or 1 % Penicillin-Streptomycin antibiotic solution (HEK293T and NIH3T3). HepG2 and Jurkat E6.1 cells were maintained in RPMI 1640 media supplemented with 10 % FBS and 1 % Penicillin-Streptomycin antibiotic solution.

### DNA and RNA transient transfections

DNA transfection in HEK293 and HEK293T cells was conducted by the well-established calcium phosphate method, using 25 μg DNA/ml of calcium phosphate solution. DNA transfection in NIH3T3 and HepG2 cells was conducted with Lipofectamine 2000 reagent following the manufacturer’s instructions (Invitrogen) and using a ratio 1μg:2μL DNA:Lipofectamine 2000. DNA transfection in ESCs was conducted with PANfect reagent following the manufacturer’s instructions (PAN biotech) and using a ratio 1μg:2μL DNA:PANfect. Jurkat E.6 cells were transfected by electroporation in 1:1 RPMI:FBS media using an exponential wave program in a Bio-Rad electroporation system and providing a single pulse of 240 V. All DNA-transfected cells were harvested 48 h post-transfection (hpt) for further analysis. siRNA/miRNA transfection for all cell lines was conducted with Lipofectamine RNAiMAX reagent following the manufacturer’s instructions (Invitrogen) and using a ratio 10pmol:2μL RNA:Lipofectamine RNAiMAX. Cells transfected with siRNAs/miRNAs were harvested 72 hpt for further analysis. For experiments requiring siRNA/miRNA and DNA transfections, DNA was transfected 24 h after siRNA/miRNA transfection. MiRNAs and siRNAs used can be found in Suppl. Materials Tables 1 and 2, respectively.

### RNA isolation, generation of cDNA libraries and RNA-seq

For RNA isolation, 5×10^6^–2×10^7^ murine ESCs were lysed in TRIZOL, and total RNA was extracted according to the manufactures’ protocol. The precipitated RNA was solved in RNAse free water. RNA quality was assessed by measuring the ratio of absorbance at 260 nm and 280 nm using a Nanodrop 2000 Spectrometer as well as by visualization of the integrity of the 28S and 18S bands on agarose gels. Total RNA was then converted into double-stranded cDNA libraries following the manufacturer’s recommendations using the Illumina TruSeq RNA Sample Preparation Kit v2. Shortly, mRNA was purified from 100 ng of total RNA using poly-T oligo-attached magnetic beads. Fragmentation was carried out using divalent cations under elevated temperature in Illumina proprietary fragmentation buffer. First-strand cDNA was synthesized using random oligonucleotides and SuperScript II. Second strand cDNA synthesis was subsequently performed using DNA Polymerase I and RNase H. Remaining overhangs were converted into blunt ends via exonuclease/polymerase activities and enzymes were removed. After adenylation of 3’ ends of DNA fragments, Illumina PE adapter oligonucleotides were ligated to prepare for hybridization. DNA fragments with ligated adapter molecules were selectively enriched using Illumina PCR primer PE1.0 and PE2.0 in a 15 cycle PCR reaction. Size-selection and purification of cDNA fragments with preferentially 200 bp in length was performed using SPRIBeads (Beckman-Coulter). Size-distribution of cDNA libraries was measured using the Agilent high sensitivity DNA assay on a Bioanalyzer 2100 system (Agilent). cDNA libraries were quantified using KAPA Library Quantification Kits (Kapa Biosystems) and were used as a template for high throughput sequencing. To this end, cluster generation was conducted on a cBot, and a 2x 100 bp paired-end run was performed on a HiSeq1500.

### RNA-seq preprocessing

Sequenced reads were aligned and quantified using STAR software (v2.7.3a)^62^ and the murine reference genome (GRCm38) from the Genome Reference Consortium. Raw counts were imported using the DESeqDataSetFromHTSeqCount function from DEseq2 (v1.26.0)^63^ and were rlog transformed according to the DEseq2 pipeline. DESeq2 was used for the calculation of normalized counts for each transcript using default parameters. All normalized transcripts with a maximum overall row mean lower than 10 were excluded, resulting in 21,685 present transcripts. RNA-seq data can be accessed under GSE163635.

### Construction of Co-expression networks analysis (CoCena)

To define differences and similarities in transcript expression patterns among the different groups, CoCena was performed based on Pearson correlation. CoCena is a network-based approach to identify clusters of genes that are co-expressed in a series of observed conditions based on data retrieved from RNA-sequencing. The tool offers a variety of functions that allow subsequent in-depth analysis of the biological context associated with the found clusters. The 10,000 most variable genes were selected as input for the analysis. A Pearson correlation coefficient cut-off of 0.915 (7,713 nodes and 410,684 edges) was chosen to construct scale-free networks.

### MiRNA enrichment analysis

Each input list of genes used for miRNA enrichment analysis consisted of CoCena gene modules found to be downregulated in a given mutant genotype as compared to the wild type counterpart (also called negative modules). Specifically, we used genes commonly downregulated in *Trim71* KO ESCs, *Lin28a* KO ESCs and double KO ESCs, derived from our own analysis (GSE163635). We additionally used genes commonly downregulated in *Trim71* KO ESCs, *Trim71* RING mutant (C12LC15A) ESCs and *Trim71* NHL mutant (R738A) ESCs, derived from a previous study which also included mRNAs enriched in TRIM71 RNA-IPs of ESCs (GSE134125)^23^. Last, we used two published datasets derived from previous studies in HCC cells^23,47^, and focused on genes significantly downregulated in *TRIM71* KO Huh7 cells (GSE134125)^23^ and genes significantly enriched in TRIM71 RNA-IPs of HepG2 cells^47^. In general, miRNA enrichment analysis was first conducted on negative modules to identify let-7 targets among the mRNAs found stabilized in the presence of TRIM71. Then, negative modules were overlapped with RNA-IP data to identify mRNAs bound and stabilized by TRIM71 (intersection). Last, miRNA enrichment analysis was conducted on the intersection to identify let-7 targets bound and stabilized by TRIM71.

miRNA enrichment analysis was performed with ShinyGo v0.61 online software^42^. This software identifies miRNA targets present in a list of genes, and returns the top X miRNAs whose targets are significantly enriched in that list based on hypergeometric distribution followed by false discovery rate (FDR) correction^42^. For this analysis, we selected a FDR cutoff of 0.05 for the top 100 enriched miRNAs. The GRIMSON (mouse) and TargetScan (human) miRNA databases were selected for the analysis of ESCs and HCC cells, respectively, as these two databases provide the enrichment for each given miRNA family. An additional analysis was then conducted to identify specific targets of individual miRNA members within each family. To this end, we used the multiMiR R package, which scans an input list of genes for predicted miRNA targets by integrating information from eight different databases (DIANA-microT, ElMMo, MicroCosm, miRanda, miRDB, PicTar, PITA and TargetScan)^64^. For miRNA network visualization, individual members of the let-7 miRNA family were depicted connected to each of their respective identified targets. Only genes predicted to be let-7 targets by at least four different databases were used for visualization. Networks were visualized using the software Cytoscape (v3.8.2).

### RNA extraction and qRT-PCR quantification

RNA was extracted from cell pellets using the Trizol-containing reagent TriFAST (peqGold) according to the manufacturer’s instructions. RNA pellets were resuspended in RNase-free water, and DNA digestion was performed prior to RNA quantification. 0.5–1 μg of RNA were reverse-transcribed to cDNA using the High Capacity cDNA Reverse Transcription Kit (Applied Biosystems) according to the manufacturer’s instructions. The cDNA was then diluted 1:5, and a relative quantification of specific genes was performed in a Bio-Rad qCycler using TaqMan probes in iTaq Universal Probes Supermix (BioRad). TaqMan probes used can be found in Suppl. Materials Table 3.

### Luciferase reporter assays

For luciferase assays, cells were co-transfected with the required psiCHECK2 dual luciferase plasmid and the specified pRK5-FLAG construct in a 1:4 ratio, and cells were harvested for further analysis 48 hpt. If miRNA overexpression or siRNA-mediated knockdown was required, miRNA/siRNA transfection was conducted 24 h before DNA transfection. psiCHECK2 plasmids contained a 3’UTR sequence with several miRNA binding sites (BS) in tandem downstream of Renilla Luciferase coding sequence, while the Firefly Luciferase, expressed under the control of a different promoter and not subjected to any altered 3’UTR regulation, was used as normalization control. To eliminate the possible 3’UTR-independent impact that each condition/construct could have on the psiCHECK2 vector backbone, a psiCHECK2 vector with no insert downstream of the Renilla sequence (Renilla-empty) was generated. Then, the Renilla-3’UTR/Firefly ratio was calculated and then normalized to the Renilla-empty/Firefly ratio calculated for the same condition. The resultant values were specified as normalized Relative Light Units (Norm. RLU). The Repression Strength (norm. RLU (FLAG-TRIM71) / norm. RLU (FLAG-Empty)) was calculated as an additional parameter representing the degree of TRIM71-mediated repression for each miRNA. Repression Strength values higher than 1 were observed upon repression of a specific miRNA, while values equal or below 1 implied that no repression was observed. Renilla and Firefly Luciferase signals were read in a MicroLumat Plus LB96V luminometer by using the Dual Luciferase Reporter Assay kit (Promega) following the manufacturers’ instructions. The 8x let-7 BS psiCHECK2 plasmid was acquired from Addgene (#20931), and the 3x miRNA BS psiCHECK2 plasmids (used in Suppl. Fig 8B) were generated by inserting artificial sequences containing three miRNA-binding sites in tandem via XhoI/NotI directional cloning.

### Protein extraction and western blotting (WB)

Cell pellets were lysed in RIPA Buffer (20 mM Tris–HCl pH 7.5, 150 mM NaCl, 1 mM Na_2_EDTA, 1 mM EGTA, 1% NP-40, 1 mM Na_3_VO_4_, 1 % sodium deoxycholate, 2.5 mM sodium pyrophosphate, 1 mM glycerophosphate) supplemented with protease inhibitors and protein lysates were pre-cleared by centrifugation and quantified using the BCA assay kit (Pierce) according to the manufacturer’s instructions. Protein lysates were then denatured by incubation with SDS Buffer (12 % glycerol, 60 mM Na2EDTA pH 8, 0.6 % SDS, 0.003 % bromophenol blue) for 10 min at 95°C and separated by PAGE-SDS in Laemmli Buffer (25 mM Tris, 192 mM glycine, 0.1 % SDS). Proteins were then wet transferred to a nitrocellulose membrane in transfer buffer (25 mM Tris–HCl pH 7.6, 192 mM glycine, 20 % methanol, 0.03 % SDS) and membranes were then blocked with 3 % BSA in 1× TBST (50 mM Tris–HCl pH 7.6, 150 mM NaCl, 0.05 % Tween-20) prior to overnight incubation at 4°C with the required primary antibodies. After washing the membrane three times with 1× TBST, they were incubated with a suitable HRP-coupled secondary antibody for 1 h at RT followed by three washing steps with 1× TBST. Membranes were developed with ECL substrate kit (Pierce) according to the manufacturer’s instructions. Antibodies used can be found in Suppl. Materials Table 4.

### Protein immunoprecipitation (IP)

For protein interaction studies, proteins were extracted on IP buffer (50 mM Tris-HCl pH 7.5, 1 mM EGTA, 1 mM EDTA, 270 mM Sucrose, 1 % Triton X-100) supplemented with phosphatase inhibitors (10 mM Glycerophosphate, 50 mM Sodium Fluoride, 5 mM Sodium Pyrophosphate, 1 mM Sodium Vanadate) and protease inhibitors. Protein lysates were pre-cleared by centrifugation at 12000 rpm and 4°C for 5 min, quantified using the BCA assay kit (Pierce), and at least 500 μg of protein were incubated with 10 μl of magnetic beads at 4°C in a rotating wheel for 5 h or overnight. A portion of each whole cell lysate was retained as input control. Sigma ANTI-FLAG M2 magnetic beads were used for the IP of Flag-tagged proteins and Protein A/G Dynabeads for the IP of Ig-tagged proteins or endogenous proteins by pre-coupling the beads to the desired primary antibody. Pre-coupling was performed in 300 μl of IP buffer with the suitable antibodies diluted 1:50 at 4 °C for 2 h in a rotating wheel. After IP, beads were washed five times with 500 μl IP buffer and bound proteins were eluted from the beads by boiling in SDS buffer, followed by PAGE-SDS separation and WB as described above. For RNase treatment after IP, washed IP fractions were incubated in 200 μl IP buffer containing 200 μg/ml RNase A at 37°C for 30 min and washed three more times with IP buffer before protein elution. Antibodies used can be found in Suppl. Materials Table 4.

### Electrophoretic mobility-shift assays (EMSA)

The generation of a recombinant TRIM71 NHL protein (FLAG-NHL) was previously described^24^. The capability of MYC-LIN28A (OriGene #TP303397) and FLAG-NHL recombinant proteins to directly bind pre-miRNAs was evaluated by EMSA. pre-let-7 and pre-miR-16 molecules (miRVana) were radioactively 5’-labeled with ^32^P isotopes by incubation with ^32^P-ƴATP and the phosphorylating enzyme T4 PNK (Roche) for 30 min at 37°C, followed by enzyme inactivation for 10 min at 65°C. ^32^P-Labeled pre-miRNAs were then purified with ProbeQuant G-50 MicroColumns (Merck) according to the manufacturer’s instructions in order to eliminate the T4 PNK enzyme and the excess of ^32^P-ƴATP. Radioactivity levels (cpm = counts per minute) of the purified samples were quantified using a liquid scintillation counter. A fixed amount of each ^32^P-labeled pre-miRNA (60 fmol) was incubated with increasing concentrations of MYC-LIN28A and/or FLAG-NHL recombinant proteins in a range of 10-1000 nM, and samples were brought to the equilibrium for 30 min at 37°C in binding buffer (10 mM MOPS pH 7.5, 50 mM KCl, 5 mM MgCl_2_, 3 mM EDTA pH 8, 30 μg/ml heparin, 5% glycerol) supplemented with 1 mM DTT and competitor yeast tRNA in a 1000-fold excess over the ^32^P-pre-miRNA to prevent unspecific binding. Control samples containing only each of the ^32^P-labeled pre-miRNAs were included to evaluate the RNA migration in the absence of protein(s). Samples were then loaded on a 10 % native PAGE gel and run for 4 h at 200 V. Gels were then dehydrated for 30-45 min at 80°C in a vacuum dryer and exposed to autoradiography films for the visualization of ^32^P-Labeled pre-miRNAs-protein complexes, detected as up-shifted bands as compared to the bands observed in control samples. Densitometry quantification of up-shifted bands was conducted with the LI-COR Image Studio Lite software and binding-saturation curves were built and adjusted by non-linear regression using GraphPad Prism7 to estimate the equilibrium dissociation constant (K_D_) of pre-miRNAs-protein complexes.

## Acknowledgements and Funding

We thank Prof. Veit Hornung and Dr. Thomas S. Ebert (Gene Center Munich, LMU) for kindly providing us with the AGO2 KO HEK293T cells. We also thank all members of our lab and the LIMES institute for general advice and discussion. LTF holds a stipend from Bayer AG and Immunosensation Cluster of Excellence. WK and JLS are funded by the Deutsche Forschungsgemeinschaft (DFG, German Research Foundation) under Germany’s Excellence Strategy EXC2151 – 390873048.

## Author Contributions

LTF and SM designed and performed all experiments, and wrote the manuscript. TU, KD, MB, KH and JLS conducted RNA-seq and bioinformatic analysis. JW conducted several experiments under the supervision of LT. WK supervised experimental design and performance, and co-wrote the manuscript.

